# FACT frees Ubp10 from nucleosome sequestration after rapid H2B deubiquitination

**DOI:** 10.64898/2026.11.06.765004

**Authors:** Raquel Merino-Urteaga, Jonas Zähringer, Melesse Nune, Cynthia Wolberger, Taekjip Ha

## Abstract

Chromatin-modifying enzymes must engage their substrate, act on it and release it, but which step limits their turnover is unclear. Using three-color single-molecule fluorescence, we directly visualized the yeast deubiquitinase Ubp10 removing ubiquitin from H2B on individual nucleosomes. Ubp10 cleaved the mark within seconds of binding, without detectable DNA unwrapping, yet it did not let go: ∼80% of nucleosomes remained bound after 45 minutes, and unmodified nucleosomes formed similarly long-lived complexes. Ubp10 turnover is therefore limited by release rather than catalysis. The histone chaperone FACT relieved this bottleneck by accelerating Ubp10-nucleosome dissociation. In living yeast, nuclear depletion of the FACT subunit Spt16 increased the slowly diffusing, chromatin-bound fraction of nucleoplasmic Ubp10. These findings identify FACT-dependent release as a likely mechanism for recycling Ubp10 on chromatin. More broadly, a nucleosome can sequester the enzyme acting on it, making enzyme release a step at which chromatin modification can be controlled.

## Introduction

Nucleosomes are the basic repeating units of eukaryotic chromatin organization, each comprising approximately 145-147 base pairs (bp) of DNA wrapped around an octamer of the core histone proteins H2A, H2B, H3, and H4^1^. The accessibility of nucleosomal DNA is dynamically regulated by histone post-translational modifications, which can alter local chromatin structure and direct the recruitment of chromatin-remodeling factors^2^. Among these modifications, monoubiquitination of histone H2B (Ub-H2B) is enriched in actively transcribed regions, where it helps define the epigenetic landscape and regulate transcription^3-5^.

Ubiquitin is conjugated to H2B during transcription initiation and throughout elongation^6, 7^, and its levels are dynamically controlled by histone deubiquitinating enzymes (DUBs). In *Saccharomyces cerevisiae*, two DUBs, Ubp8 and Ubp10, act on distinct pools of Ub-H2B^8-11^. Ubp8 functions within the SAGA complex^12-14^ near promoters and transcription start sites, where it assists the transition of RNA polymerase II into productive elongation^8^. Ubp10, the yeast counterpart of human USP36^15^, is a monomeric enzyme that targets mid-coding regions and subtelomeres, regulating gene expression in euchromatin and facilitating telomeric gene silencing^10, 11, 16-18^.

Earlier studies reported that Ubp10 deubiquitinates Ub-H2B within the free H2A/H2B heterodimer complex, whereas its activity is much weaker on a ubiquitinated nucleosome^17, 19^. The intrinsically disordered N-terminus of Ubp10 forms extensive contacts with the free dimer that are diminished or reorganized in the nucleosomal context, providing a possible structural explanation for this difference in activity^20^. These observations suggested that ubiquitinated H2B may be an intrinsically poor substrate for Ubp10 once packaged into a nucleosome, but the kinetic basis for this weak activity is undefined.

The histone chaperone FACT (Facilitates Chromatin Transcription) strongly stimulates Ubp10 activity on nucleosomes, as shown in bulk experiments^19^. FACT, comprising Spt16-Pob3 together with the DNA-binding protein Nhp6^21-24^, reorganizes nucleosomes during RNA polymerase II transcription^23, 25-28^ and promotes local histone recycling^29^. Its function is also closely linked to H2B ubiquitination during elongation, including suppression of cryptic and antisense transcription^4, 30, 31^. How these activities translate into faster Ubp10 turnover, and which kinetic bottleneck FACT relieves, remains unknown.

Single-molecule approaches can resolve transient and heterogeneous chromatin transactions that are obscured by ensemble averaging. They have provided mechanistic insight into nucleosome interactions with polymerases^32, 33^, transcription factors^34, 35^, chromatin remodelers^36, 37^, and histone chaperones^33^. Moreover, these methods have been applied to monitor ubiquitin conjugation^38^ and ubiquitin-dependent protein processing^39, 40^, but they have not yet been used to follow the removal of a ubiquitin mark from chromatin. Direct visualization of a deubiquitinase such as Ubp10 acting on an individual intact nucleosome substrate is needed to identify the kinetic steps that limit its activity on chromatin and how FACT overcomes this constraint.

Here, we used a three-color single-molecule nucleosome assay to monitor Ubp10-catalyzed ubiquitin removal, nucleosomal DNA wrapping, and substrate residence. The enzyme cleaved the ubiquitin mark within seconds of binding without detectable DNA unwrapping, yet remained associated with the nucleosome for more than 45 minutes. FACT accelerated dissociation of this long-lived complex *in vitro*, and nuclear depletion of the FACT subunit Spt16 raised the slowly diffusing fraction of nucleoplasmic Ubp10 in living cells, consistent with greater chromatin association. Together with previous evidence of FACT-dependent stimulation in bulk activity assays when nucleosomes are present in excess, these findings establish a mechanism in which FACT releases and recycles nucleosome-bound Ubp10. More broadly, our results identify nucleosome-mediated sequestration as a potential constraint on the turnover of chromatin-associated enzymes.

## Results

### Ubp10 deubiquitinates single nucleosomes without detectable DNA unwrapping

We adapted a three-color ubiquitinated nucleosome substrate, previously developed to monitor FACT-mediated DNA unwrapping and H2A/H2B dimer eviction^20^, to measure Ubp10 deubiquitination on individual surface-immobilized nucleosomes. Nucleosomes containing *S. cerevisiae* core histones were assembled on DNA carrying the Widom 601 sequence extended at each end by a 43-base pair (bp) DNA linker. Cy3 and Cy5 were positioned at superhelical location +1 and on the linker DNA, respectively, such that a fully wrapped nucleosome exhibited high FRET efficiency (Fig. 1a; sequence in Supplementary Table S1). Cy7-labeled ubiquitin was enzymatically conjugated to H2B via a native isopeptide bond (see Methods). Because each nucleosome contains two copies of H2B and ubiquitin dye-labeling efficiency was approximately 50%, the Cy7 intensity distribution contained populations corresponding to nucleosomes with zero, one, or two Cy7-labeled ubiquitins (Extended Data Fig. 1a). This substrate therefore allowed simultaneous monitoring of H2B deubiquitination through Cy7 loss and nucleosomal DNA wrapping through Cy3-Cy5 FRET.

**Figure 1:**
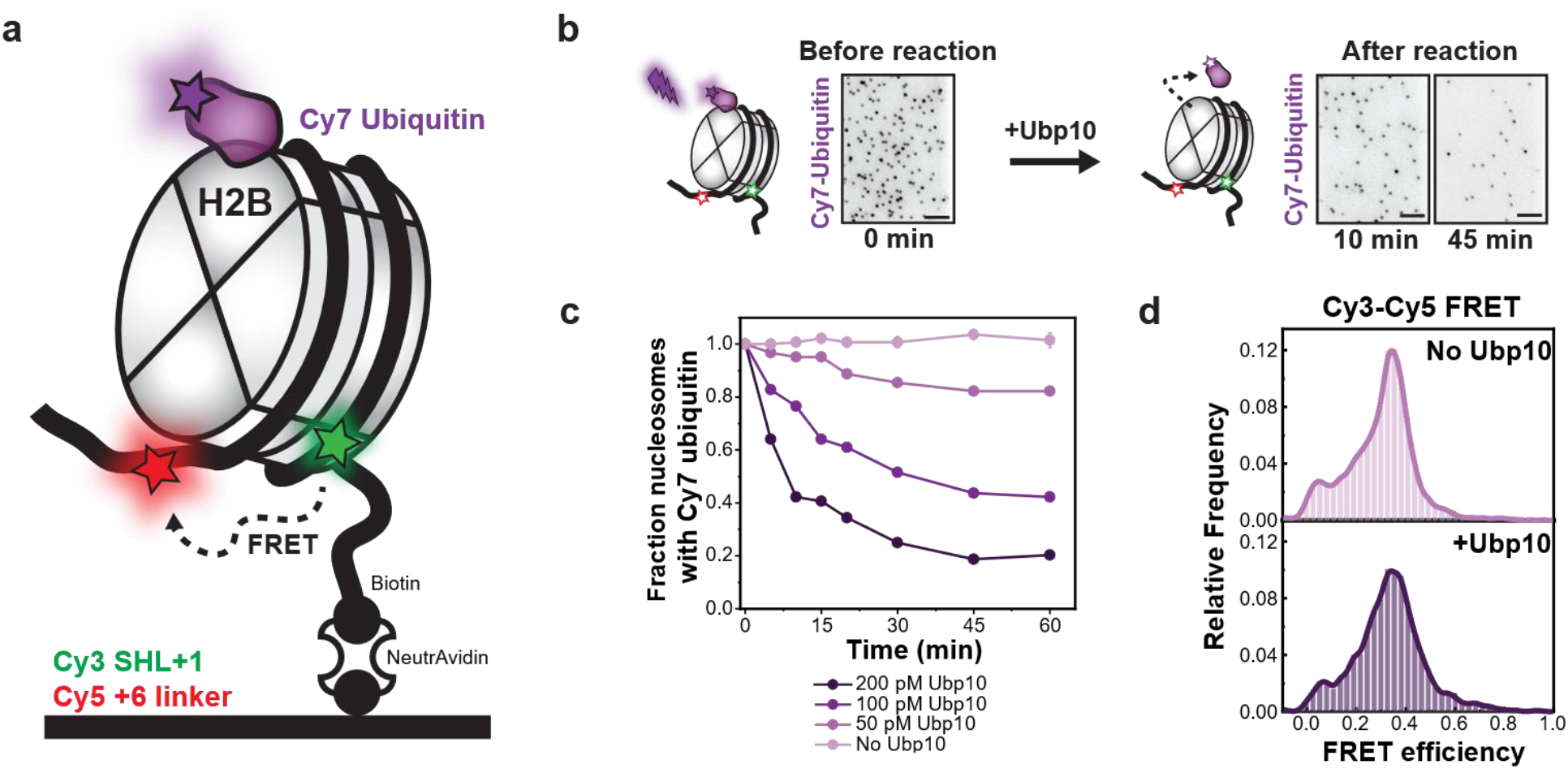
Single-molecule platform to study H2B deubiquitination on nucleosomes. **a,**Schematic of the three-color nucleosome construct designed to simultaneously observe DNA dynamics and ubiquitin-H2B cleavage. **b,** Deubiquitination detected through the loss of Cy7 spots colocalized with Cy3/Cy5 nucleosomes. Representative Cy7-ubiquitin fields show that the number of Cy7 spots decreases over time after Ubp10 addition (0, 10, and 45 min; scale bar, 10 µm). **c,** Time-resolved ubiquitin cleavage on surface-tethered Ub-H2B nucleosomes as a function of Ubp10 concentration. **d,** Stable Cy3-Cy5 FRET efficiency upon Ubp10 binding demonstrates intact nucleosomal DNA wrapping post-cleavage.

Addition of Ubp10 caused a rapid decrease in Cy7-positive nucleosomes (Fig. 1b and Extended Data Fig. 1b), with ∼60% deubiquitinated within 10 minutes at 200 pM Ubp10 (Fig. 1c). No comparable Cy7 loss was observed with catalytically inactive Ubp10(C371A) (Extended Data Fig. 1c). Neither Ubp10 binding nor subsequent deubiquitination produced any change in Cy3-Cy5 FRET (Fig. 1d). Thus, ubiquitin cleavage occurred without a detectable change in nucleosomal DNA wrapping under the conditions tested.

### Ubp10 cleaves nucleosomal ubiquitin within seconds of binding

Having established that Ubp10 deubiquitinates intact, surface-immobilized nucleosomes, we next measured the timing of ubiquitin cleavage relative to initial nucleosome engagement. To do so, we reversed the assay geometry by tethering Ubp10-mGold2t to the surface through a biotinylated anti-GFP antibody, an approach previously validated for specific recognition of mGold2t-tagged proteins^41^ (Fig. 2a). Fluorescently labeled Ub-nucleosomes were then introduced and their recruitment monitored by Cy5 fluorescence. Control surfaces lacking Ubp10 showed minimal nonspecific adsorption, indicating that approximately 80% of captured nucleosomes interacted specifically with Ubp10 (Extended Data Fig. 2a-c).

**Figure 2:**
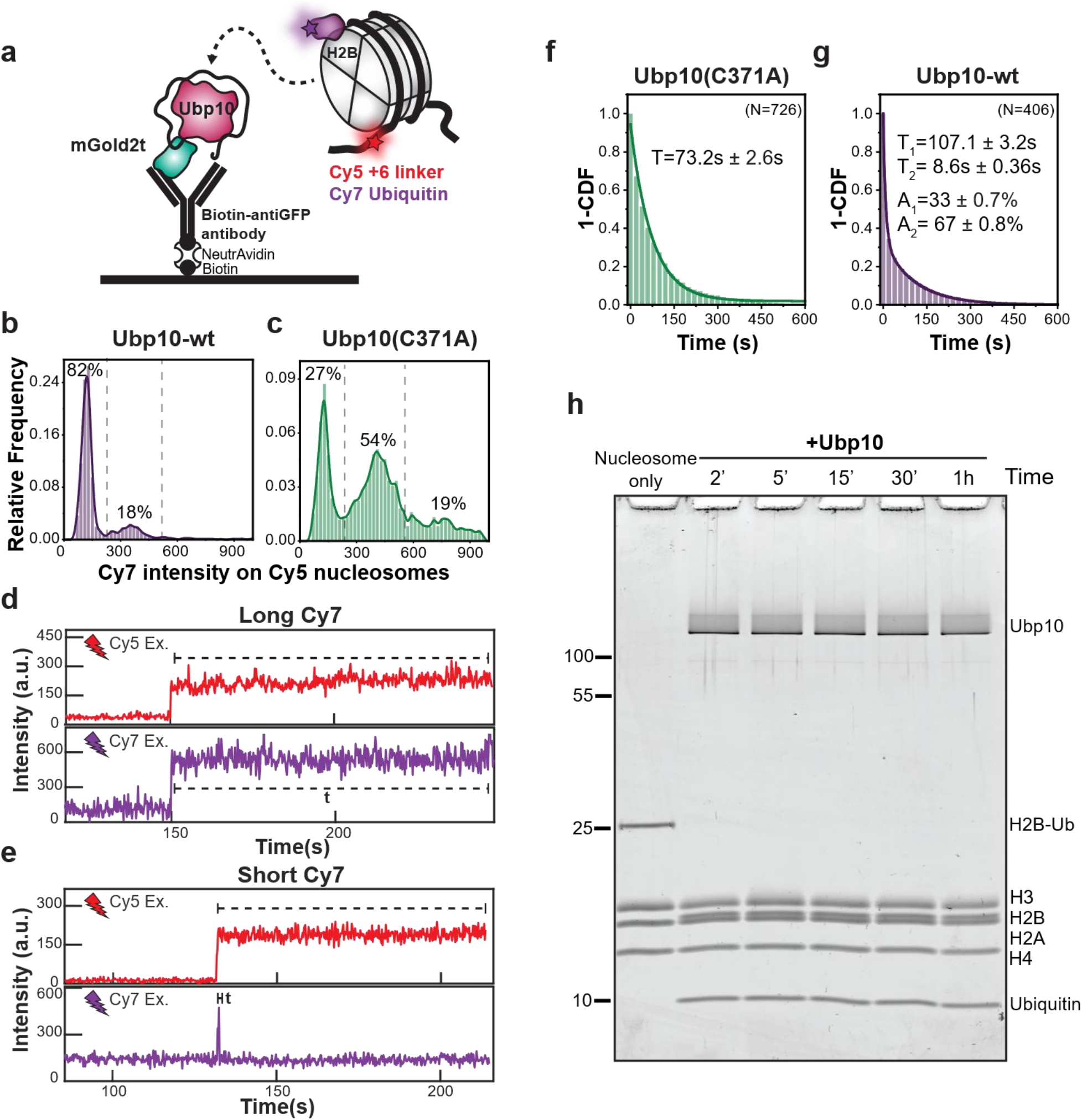
Real-time single-molecule enzyme kinetics on ubiquitinated nucleosomes by surface-tethered Ubp10. **a,**Experimental design illustrating fluorescent nucleosome binding to Ubp10-mGold2t immobilized on the surface using a biotinylated anti-GFP antibody. **b,c,** Cy7 fluorescence intensity histograms of nucleosomes bound to wild-type Ubp10-mGold2t (Ubp10-wt) (**b**); or the catalytically inactive mutant Ubp10(C371A)-mGold2t (**c**). The three populations correspond to nucleosomes carrying zero, one, or two Cy7-labeled ubiquitins (dashed lines). Nucleosomes bound to the inactive mutant retain all three populations, whereas binding to active Ubp10 depletes the one- and two-Cy7 populations and increases the zero-intensity background population, confirming enzymatic cleavage. **d,e,** Representative single-molecule trajectories depicting nucleosome landing events, marked by simultaneous Cy5 and Cy7 signal onset. Two distinct Cy7 dwell times were observed: (**d**) long-lived and (**e**) short-lived. **f,g** Survival probability curves (1-CDF) of Cy7 lifetimes for nucleosomes interacting with (**f**) Ubp10(C371A) or (**g**) Ubp10-wt. The photobleaching lifetime of Cy7 was determined using the inactive mutant to be ∼73 s (**f**). Ubp10-wt exhibited double decay kinetics (**g**): a fast phase (T2=∼9 s) corresponding to rapid enzymatic cleavage, and a slow phase (T1=∼100 s) representing non-specifically bound species and photobleaching. **h**, Bulk DUB activity assay at equimolar enzyme-to-substrate concentrations demonstrating complete deubiquitination of H2B on nucleosomes after 2 min of Ubp10 incubation.

The Cy7 intensity distribution of Ubp10-bound nucleosomes contained two populations: a major population with background-level Cy7 signal (∼82%), consistent with loss of Cy7-labeled ubiquitin, and a minor population retaining one Cy7-labeled ubiquitin (∼18%) (Fig. 2b). Conversely, nucleosomes captured by catalytically inactive Ubp10(C371A)-mGold2t retained the baseline three-population Cy7 distribution, with no detectable deubiquitination (Fig. 2c and Extended Data Fig. 1a). Thus, surface-tethered wild-type Ubp10 deubiquitinated the nucleosomes it captured, whereas the catalytically inactive mutant did not.

To determine how rapidly this reaction occurred after nucleosome capture, we monitored the duration of Cy7 fluorescence after the coincident appearance of Cy5 (nucleosome) and Cy7 (ubiquitin) signals on the Ubp10-functionalized surface (Fig. 2d and 2e). With catalytically inactive Ubp10(C371A) and on NeutrAvidin control surfaces, Cy7 signals persisted for ∼70–110 seconds, establishing the photobleaching baseline (Fig. 2d, 2f, and Extended Data Fig. 2d-f). By contrast, the Cy7 lifetime distribution measured with wild-type Ubp10-mGold2t was best described by a double-exponential decay (Fig. 2g). The fast component (Fig. 2e), accounting for ∼67% of the fitted amplitude, had a lifetime of less than nine seconds and was therefore attributed to Ubp10-catalyzed ubiquitin cleavage. The slow component (∼100 seconds) matched the lifetime of the photobleaching controls (Fig. 2d, 2f, and Extended Data Fig. 2d-f) and likely reflects the small fraction of nucleosomes adsorbed nonspecifically to the surface.

As an orthogonal test, we measured bulk DUB kinetics at an equimolar enzyme-to-substrate ratio using nucleosomes reconstituted with commercially sourced human octamers carrying site-specifically monoubiquitinated H2BK120 (H2BK120ub1; Extended Data Fig. 3a), equivalent to the conserved yeast H2BK123 site. Ubp10 completely removed ubiquitin from these nucleosomes within two minutes (Fig. 2h and Extended Data Fig. 3b), consistent with the rapid cleavage observed in the single-molecule assay with enzymatically ubiquitinated nucleosomes. Together, these measurements show that Ubp10 cleaves nucleosomal ubiquitin within seconds of binding.

### Slow nucleosome release restricts Ubp10 turnover

Previous bulk assays reported weak Ubp10 activity on nucleosomes at low enzyme-to-substrate ratios^19^, whereas our single-molecule and equimolar bulk measurements showed rapid deubiquitination upon nucleosome binding. We therefore asked whether slow nucleosome release, rather than ubiquitin cleavage, limits Ubp10 turnover at low enzyme concentrations.

Although the Cy7-ubiquitin signal disappeared within seconds, the Cy5-nucleosome signal persisted in the same single-molecule trajectories (Fig. 2e), indicating that ubiquitin removal was not immediately followed by nucleosome dissociation. To quantify this, we washed away free nucleosomes from the TIRF channel and tracked the number of Cy5-labeled nucleosomes remaining on the Ubp10-functionalized surface over time. Approximately 80% stayed bound to Ubp10 after 45 minutes (Fig. 3a). Unmodified nucleosomes tethered to the same surface yielded comparable binding efficiency and complex stability (Extended Data Fig. 4a and 4b). Thus, long-lived binding is an intrinsic property of the Ubp10-nucleosome interaction rather than a state induced specifically by ubiquitin cleavage.

**Figure 3:**
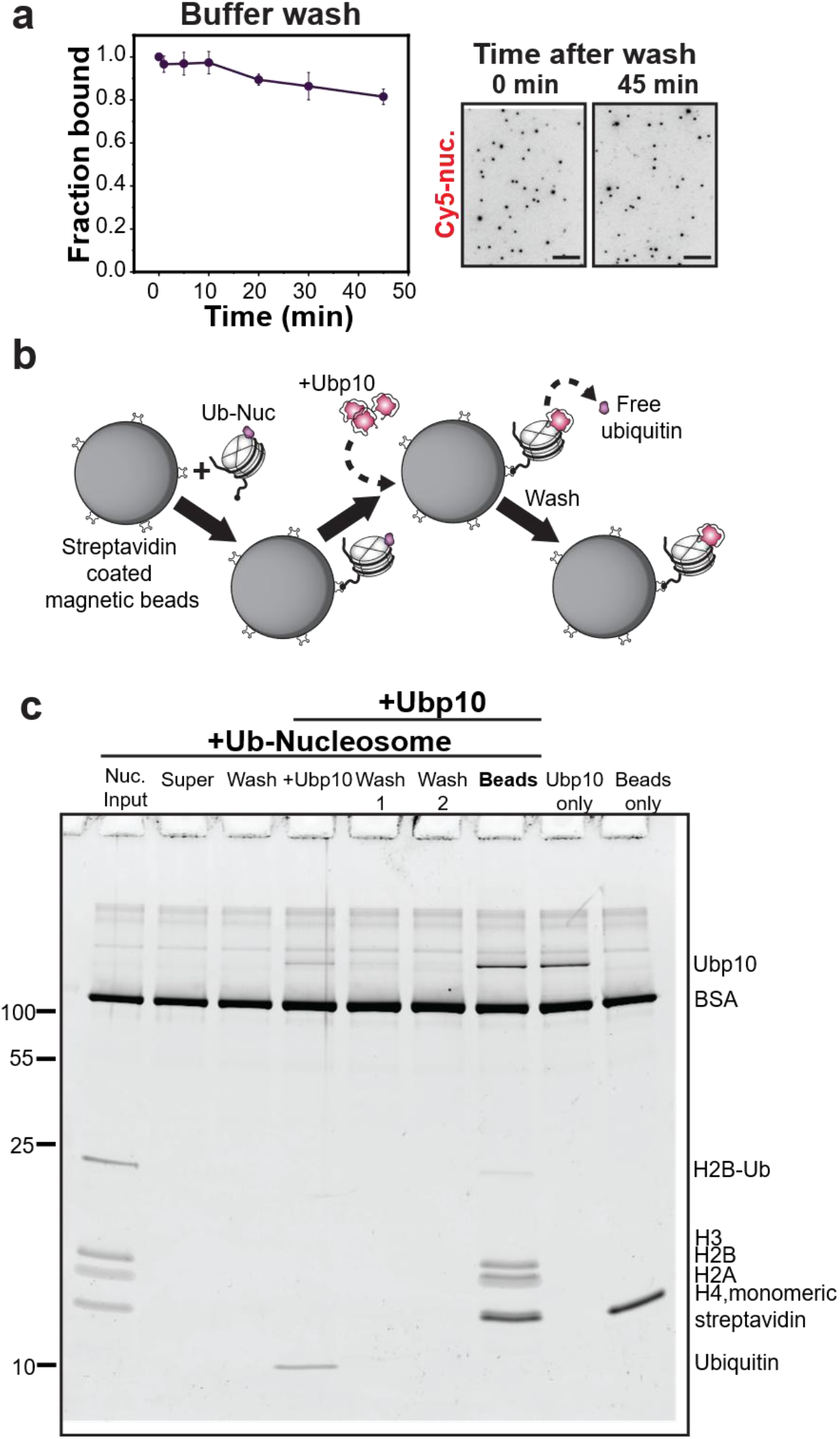
Ubp10 remains stably bound to nucleosomes after ubiquitin cleavage. **a**, Time-course tracking of Cy5-nucleosomes showing retention on surface-tethered Ubp10 after washing away unbound substrate from the flow channel (scale bar, 10 µm). Error bars are SEM (n=3). **b**, Schematic of the on-bead bulk deubiquitination assay. Biotinylated ubiquitinated nucleosomes (265 nM) were immobilized onto streptavidin-coated magnetic beads and incubated with Ubp10 (265 nM). Following wash steps, stably bound complexes on beads were eluted by SDS denaturation. **c**, SDS-PAGE gel showing co-elution of Ubp10 alongside histone proteins upon bead denaturation, confirming stable complex formation.

**Figure 4:**
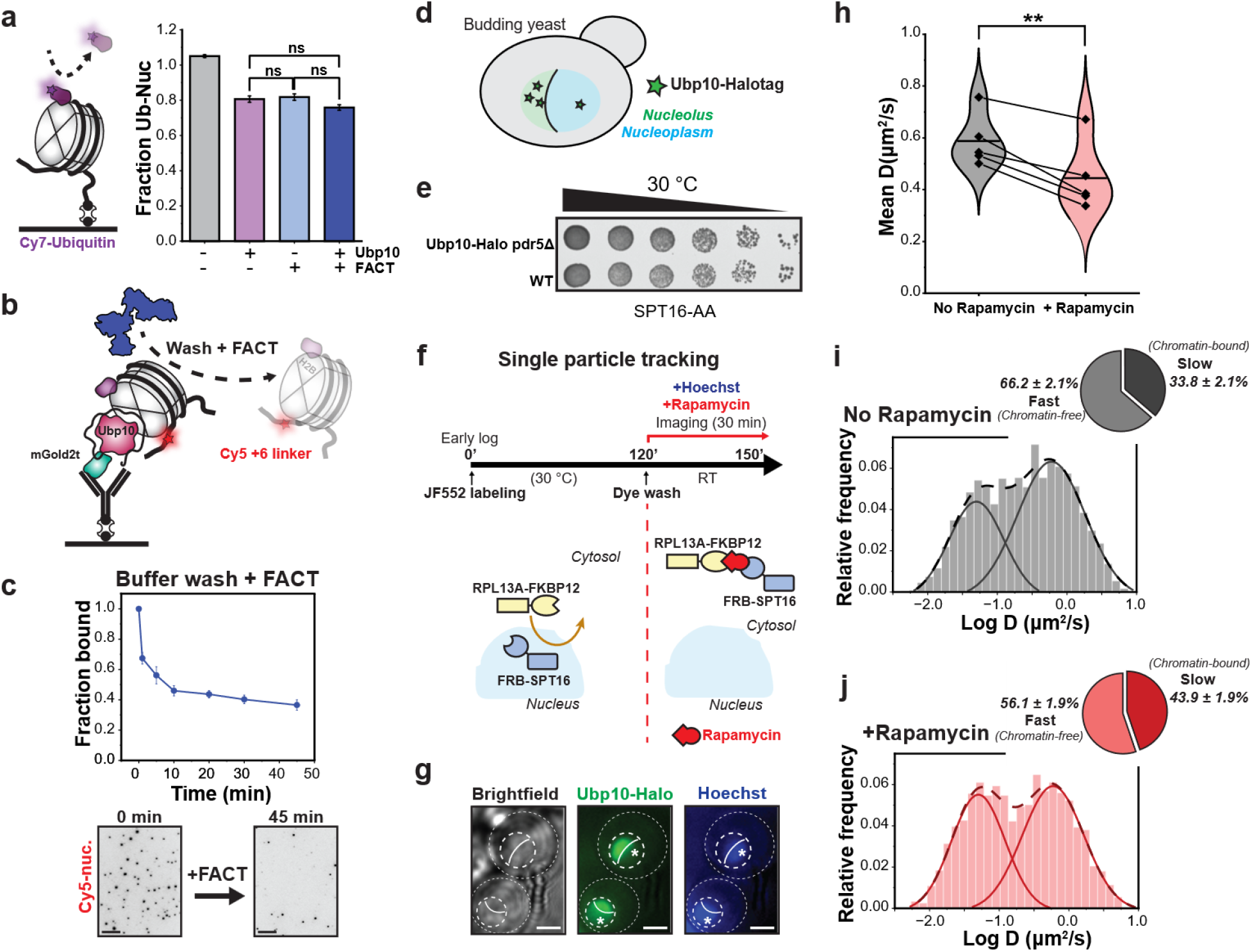
The FACT complex promotes Ubp10 dissociation from chromatin in vitro and in cells. **a**, Biotinylated, fluorescently labeled Ub-nucleosomes were immobilized on NeutrAvidin-coated slides. Fraction of Ub-nucleosomes (Cy7 spots) remaining was measured after 45 min of incubation with Ubp10 (10 pM) in the presence or absence of FACT (1 μM Spt16-Pob3 + 2 μM Nhp6a). FACT supplementation failed to stimulate Ubp10 activity. Error bars are SEM. Statistical differences were determined by unpaired t-test (n=3, ns = p > 0.05). **b**, Experimental scheme for FACT-mediated wash-off assays on Ub-nucleosomes captured by surface-tethered Ubp10. **c**, Time course of nucleosome release from surface-tethered Ubp10 after addition of FACT, quantified as the fraction of Cy5-nucleosome spots remaining bound. Representative fields at 0 and 45 min are shown below (n=3; scale bar, 10 µm). **d**, Schematic of a budding yeast cell expressing Ubp10-HaloTag, illustrating its subnuclear partitioning between the nucleolus and nucleoplasm. **e**, Fivefold serial dilution growth assays comparing growth of Ubp10-HaloTag and untagged control strains in an Spt16-AA, pdr5Δ background on YPD agar at 30 °C for 2 days. **f**, Schematic of live-cell single-particle tracking (spTracking) setup. Early log-phase Spt16-AA yeast cells expressing Ubp10-HaloTag were labeled with 50 nM JF552 for two hours at 30 °C, washed, and immobilized on concanavalin A-coated coverslips. Hoechst dye was added for nucleoplasm staining. Time-course imaging was performed over 30 minutes of rapamycin treatment (nuclear Spt16 depletion). **g**, Subnuclear localization of JF552-Ubp10 relative to Hoechst-stained chromatin using HILO illumination, illustrating spatial masking used to segment nucleoplasmic trajectories (scale bar, 2 µm). **h**, Mean apparent Ubp10 diffusion coefficient decreases following rapamycin treatment. Lines connect the same cells imaged before and after rapamycin addition. Error bars are SEM. Statistical differences were determined by paired t-test (n=5, ** = p < 0.01). **i,j**, Two-state Gaussian fits of log(D) distributions at (**i**) no rapamycin vs. (**j**) + rapamycin (movies acquired 15–30 min after addition; see Methods), highlighting an increased Ubp10 chromatin-bound fraction upon Spt16 nuclear depletion (n=5).

Using a complementary approach, we immobilized biotinylated ubiquitinated nucleosomes onto streptavidin-coated magnetic beads (Fig. 3b and Extended Data Fig. 4c). Addition of Ubp10 resulted in the release of free ubiquitin, confirming its catalytic activity on the bead surface. After washing away unbound protein, bead-associated material was recovered by denaturation in SDS sample buffer. Most of the Ubp10 was co-eluted with the histones (Fig. 3b and 3c). In contrast, beads lacking tethered nucleosomes retained only trace amounts of non-specifically bound Ubp10 (Extended Data Fig. 4d). Together with the single-molecule residence-time measurements, these results demonstrate that Ubp10 forms a long-lived complex with nucleosomes. Thus, rapid ubiquitin cleavage is followed by slow nucleosome release, restricting Ubp10 availability for subsequent catalytic cycles.

### FACT accelerates dissociation of the Ubp10-nucleosome complex

Prior work showed that FACT stimulates Ubp10-dependent deubiquitination of nucleosomes^19^, but the underlying mechanism remained unclear. We first asked whether FACT increased the extent of deubiquitination of surface-immobilized nucleosomes (Fig. 4a). At the Ubp10 concentrations used above, deubiquitination was too rapid to resolve any additional effect of FACT, so we reduced the nominal Ubp10 concentration to 10 pM, at which approximately 20% of the substrate was deubiquitinated over 45 minutes. FACT did not significantly increase the extent of deubiquitination under these conditions: the loss of Cy7 signal was similar between Ubp10 alone, FACT alone (which evicts Ub-H2B/H2A dimers^20^) and both combined (Fig. 4a). Because deubiquitination at 10 pM was much slower than at higher Ubp10 concentrations, the reaction was likely limited by Ubp10 binding to nucleosomes rather than by enzyme turnover, a regime in which FACT-dependent release would not be expected to increase activity.

Because this endpoint assay did not distinguish substrate engagement, catalysis, and enzyme release, we next tested directly whether FACT acts on the long-lived Ubp10-nucleosome complex. Nucleosomes were first allowed to bind surface-tethered Ubp10, after which the resulting complexes were washed with buffer containing FACT (Fig. 4b). FACT markedly accelerated nucleosome dissociation: more than 30% of bound nucleosomes were released within the first minute, and approximately 60% had dissociated by 10 minutes (Fig. 4c). After 45 minutes, only about 30% of the initial population remained, compared with 80% in the absence of FACT. Thus, FACT directly accelerates dissociation of the long-lived Ubp10-nucleosome complex, providing a mechanism by which FACT could restore Ubp10 turnover.

### Nuclear FACT depletion increases Ubp10 chromatin residence in living cells

To evaluate how the FACT complex shapes Ubp10 dynamics *in vivo*, we used single-particle tracking (spTracking) in a budding yeast strain in which HaloTag was fused to endogenous Ubp10 (Fig. 4d) and the FACT subunit Spt16 can be conditionally depleted from the nucleus by Anchor-Away^42^ (Fig. 4f). In-gel fluorescence of the covalently bound HaloTag ligand and anti-HaloTag immunoblotting detected full-length Ubp10-HaloTag, with no prominent lower-molecular-weight products (Extended Data Fig. 5a and 5b).

**Figure 5:**
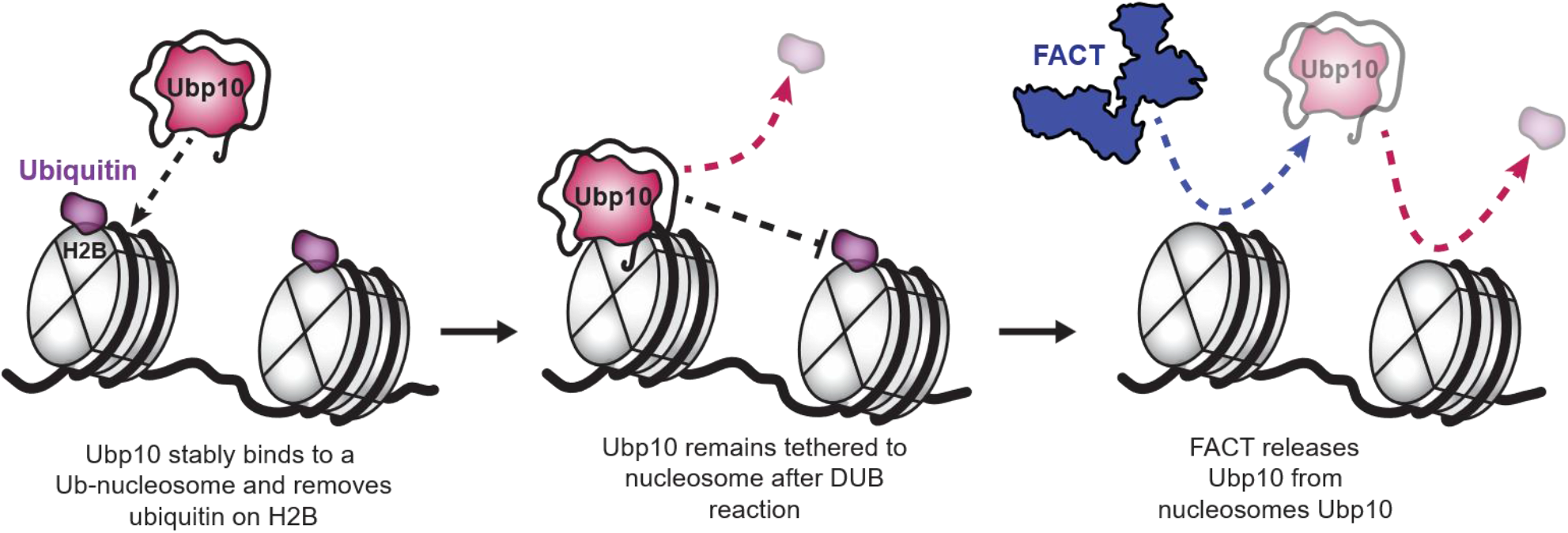
Proposed model for FACT-facilitated Ubp10 turnover on nucleosomes. Left, Ubp10 binds an H2B-ubiquitinated nucleosome and removes ubiquitin from H2B within seconds of binding. Middle, after cleavage Ubp10 remains bound to the deubiquitinated nucleosome, so neighboring ubiquitinated nucleosomes are not targeted. Right, binding of FACT to the Ubp10-nucleosome complex releases Ubp10, which can then bind and deubiquitinate another nucleosome.

Growth of this strain was comparable to that of the corresponding untagged Anchor-Away control, and the fusion did not detectably alter baseline Spt16 or histone H3 levels (Fig. 4e, Extended Data Fig. 5c and 5d). Chromatin fractionation established that 30 minutes of rapamycin treatment produced ∼75% Spt16 depletion while leaving histone H3 largely unchanged (Extended Data Fig. 6).

For spTracking, early log-phase cells were labeled to saturation with the JF552-HaloTag ligand and immobilized (Fig. 4f). Because Ubp10 is enriched in the nucleolus but also acts on H2B in the nucleoplasm, we used Hoechst staining to distinguish between the two compartments. Most JF552-Ubp10 fluorescence localized to a Hoechst-poor region within the nucleus, consistent with its expected enrichment. We therefore restricted trajectory analysis to the Hoechst-positive nucleoplasm (Fig. 4g).

Apparent diffusion coefficients (*D)* of individual trajectories were measured at 10-ms temporal resolution, and the log(*D*) distribution was fitted with a two-state Gaussian model. The slow population had an apparent diffusion coefficient of *D_slow_* = 0.05 µm² s⁻¹, consistent with chromatin association, whereas the fast population had an approximately tenfold higher coefficient of *D*_fast_ = 0.59 µm^2^s^−1^ (Fig. 4i). After 15-30 min of rapamycin treatment, the mean apparent diffusion coefficient decreased (Fig. 4h), accompanied by an increase in the slow fraction from 34% to 44% (Fig. 4i, j). To determine whether rapamycin treatment alone could account for this shift, we performed spTracking in the Ubp10-HaloTag parental control strain lacking the Spt16-FRB fusion (Extended Data Fig. 5b and 5d). There, rapamycin modestly decreased the slow fraction and increased mean Ubp10 diffusivity (Extended Data Fig. 7), changes opposite in direction to those observed after Spt16 Anchor-Away. Thus, the increase in the slow Ubp10 population is driven by nuclear Spt16 depletion rather than by rapamycin itself. Overall, these findings support a role for FACT in limiting Ubp10 residence on chromatin *in vivo*, consistent with FACT-dependent dissociation of the Ubp10-nucleosome complex observed *in vitro*.

## Discussion

Ubp10 cleaves ubiquitin far more efficiently from free Ub-H2B/H2A heterodimers than from ubiquitinated nucleosomes at low enzyme-to-substrate ratios in bulk assays^17, 19^. These observations suggested that the nucleosome is a poor catalytic substrate for Ubp10. By contrast, our single-molecule measurements show that Ubp10 cleaves nucleosomal ubiquitin within seconds of binding. These results resolve the apparent discrepancy by distinguishing catalysis from nucleosome release. Although cleavage was rapid, approximately 80% of nucleosomes remained associated with Ubp10 after 45 minutes. Unmodified nucleosomes formed similarly stable complexes, consistent with previous electrophoretic mobility shift assays showing comparable Ubp10 affinity for ubiquitinated and unmodified nucleosomes^19^. Thus, weak activity under limiting-enzyme conditions does not reflect slow ubiquitin cleavage. Instead, slow nucleosome release after rapid catalysis restricts Ubp10 availability for subsequent catalytic cycles.

The previously reported preference of Ubp10 for free Ub-H2B/H2A heterodimers was thought to have a structural basis. Cross-linking mass spectrometry detected fewer contacts between Ubp10 and H2A/H2B in nucleosomes than in free heterodimers, along with new contacts formed on H3 and H4^20^. AlphaLink2 models^43, 44^ based on these data predicted that the heterodimer-bound orientation of Ubp10 would clash with nucleosomal DNA and with histones H3 and H4, suggesting that the enzyme must engage the nucleosome differently, in a configuration of unknown catalytic competence^20^. Our measurements show that Ubp10 is nevertheless active in this alternative binding mode. Thus, the distinct pattern of Ubp10-histone contacts on nucleosomes is compatible with rapid catalysis and, as our data show, does not require detectable DNA unwrapping.

Ubp10 acts on Ub-H2B nucleosomes within actively transcribed gene bodies, particularly within longer genes marked by H3K79me3^18, 19^. Sustained deubiquitination in these regions would therefore require factors that release nucleosome-bound Ubp10 and permit additional catalytic cycles. FACT is a strong candidate for this role. Its subunits Spt16 and Pob3 have been detected together with core histones in Ubp10 interaction proteomics^45, 46^, and FACT-deficient strains accumulate Ub-H2B^19^. Combined inactivation of FACT and Ubp10 also delays recovery from replication stress and increases cryptic transcription linked to Ub-H2B accumulation^47^. Previous bulk assays showed that FACT strongly stimulates Ubp10-dependent deubiquitination when nucleosomal substrate is present in excess, suggesting that FACT promotes enzymatic turnover. This effect was specific to Ubp10, as FACT did not comparably enhance Ubp8 activity^13, 19^. However, the molecular step underlying FACT-dependent stimulation remained unknown.

Our single-molecule measurements identify accelerated dissociation of the Ubp10-nucleosome complex as the molecular basis for FACT-dependent stimulation. In the present surface-immobilized nucleosome assay, FACT did not detectably increase deubiquitination at low Ubp10 concentration, possibly because substrate engagement rather than enzyme release limited the reaction. By contrast, FACT strongly accelerated dissociation of nucleosomes already bound to Ubp10. Together, the earlier bulk results and our direct measurements show that FACT promotes Ubp10 recycling by releasing the enzyme from a long-lived nucleosome complex and enabling subsequent catalytic cycles.

Live-cell spTracking provided independent support for this model. We restricted the analysis to nucleoplasmic Ubp10, excluding the major nucleolar pool, where it primarily acts on RNA polymerase I (Rpa190) to sustain rRNA production and ribosome biogenesis^16, 46^. The slow population had an apparent diffusion coefficient of 0.05 µm^2^s^−1^, consistent with chromatin association and comparable to values reported for other chromatin-interacting proteins^48-50^. Nuclear depletion of Spt16 increased the slow fraction whereas rapamycin treatment shifted the distribution in the opposite direction in the parental control strain. These observations extend our *in vitro* mechanism to the living nucleus, nominating FACT as a physiological regulator of Ubp10’s residence on chromatin. The untreated slow fraction was lower in the Spt16-FRB strain than in the parental strain (34% versus 52%; Fig. 4i and Extended Data Fig. 7a), and the tagged strain grew more slowly (Extended Data Fig. 5d). This is consistent with the FRB domain, fused adjacent to the acidic Spt16 region that binds H2A/H2B^51, 52^, partially altering FACT function before rapamycin addition. All comparisons were therefore made within each strain, before and after depletion.

Whether the persistent Ubp10-nucleosome interaction extends to polynucleosomes or longer chromatin fibers remains unknown. A previous pulldown study found that Ubp10 bound poorly to reconstituted 3-kb chromatin unless it was co-assembled with the SIR complex^17^. Those experiments were conducted in 10 mM magnesium acetate, conditions expected to favor chromatin compaction^53^, likely yielding a substantially less accessible substrate than the mononucleosomes used here. In subtelomeric heterochromatin, Ubp10 may instead depend on interactions between its intrinsically disordered N-terminal region and the SIR complex for efficient recruitment^10, 16^. Further experiments using defined nucleosome arrays under different compaction conditions will be required to determine how chromatin organization influences Ubp10 binding, release, and FACT dependence.

The slowly diffusing nucleoplasmic population is unlikely to consist exclusively of Ubp10 bound to Ub-H2B nucleosomes on its own. As noted above, Ubp10 can associate with the SIR complex to target condensed heterochromatin^10, 16^. Additionally, it may engage in other nucleoplasmic transactions, such as deubiquitinating PCNA at replication forks^54^. Because PCNA ubiquitination is low under the unperturbed conditions used here^55^, this contribution is likely small, although other FACT-independent Ubp10 complexes cannot be excluded. Accordingly, the spTracking measurements show that Spt16 depletion shifts Ubp10 toward slowly diffusing nucleoplasmic complexes but do not establish which substrate or interaction accounts for this change.

Taken together, our study separates two kinetically distinct steps in Ubp10-mediated nucleosomal H2B deubiquitination. After binding an intact nucleosome, Ubp10 rapidly cleaves ubiquitin but remains bound in a long-lived complex with the nucleosome that restricts further turnover. FACT accelerates dissociation of this complex *in vitro*, while nuclear depletion of Spt16 increases the slowly diffusing Ubp10 population in living cells. We therefore propose that FACT binding and nucleosome reorganization displace nucleosome-bound Ubp10, recycling the enzyme in chromatin (Fig. 5). Other histone chaperones or chromatin-associated factors may also promote Ubp10 release, but our findings identify FACT as a regulator of this previously unrecognized kinetic step.

FACT may also stimulate Ubp10 through a second mechanism. By fully releasing Ub-H2B/H2A heterodimers during nucleosome reorganization it could supply excellent, rapidly processed substrates for deubiquitination. This, however, is likely uncommon, because FACT typically keeps the displaced H2A/H2B dimer tethered through the C-terminal region of Spt16 rather than releasing it, preserving chromatin stability^25, 26^, and the eviction that does occur is further reduced roughly twofold on Ub-nucleosomes^20^. Thus, FACT may sustain Ubp10 activity on chromatin mainly by releasing the enzyme from a long-lived nucleosome complex, rather than by generating free dimer substrates. FACT-dependent release may represent a more general mode of recycling proteins that form long-lived complexes on nucleic acids. Consistent with this possibility, FACT has been reported to convert Cas9 from single- to multi-turnover behavior by promoting its dissociation from DNA substrates^56^.

## Extended Data Figures

**Extended Data Figure 1:**
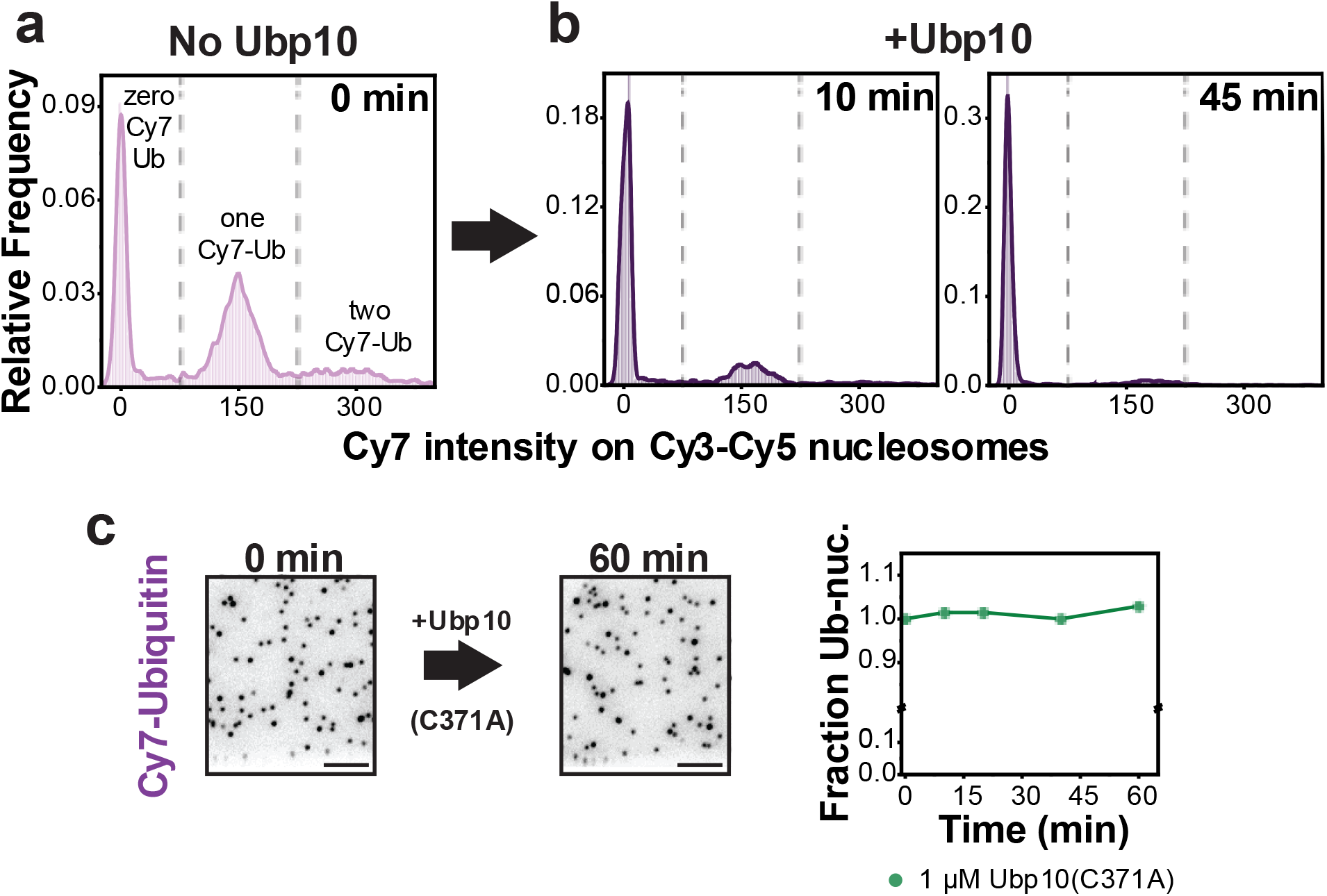
Deubiquitinase activity of wild-type Ubp10 and catalytically dead Ubp10(C371A) on nucleosomes. **a**, Cy7 fluorescence intensity histograms of surface-immobilized ubiquitinated nucleosomes. Three distinct populations are resolved, corresponding to nucleosomes bearing zero, one, or two Cy7-ubiquitin moieties. **b**, Cy7 intensity histograms after wild-type Ubp10 addition (10 and 45 min), displaying a time-dependent accumulation of non-ubiquitinated nucleosomes. **c**, Time-course quantification of the ubiquitinated nucleosome fraction upon incubation with catalytically dead Ubp10(C371A), demonstrating no detectable ubiquitin cleavage over time (scale bar, 10 µm).

**Extended Data Figure 2:**
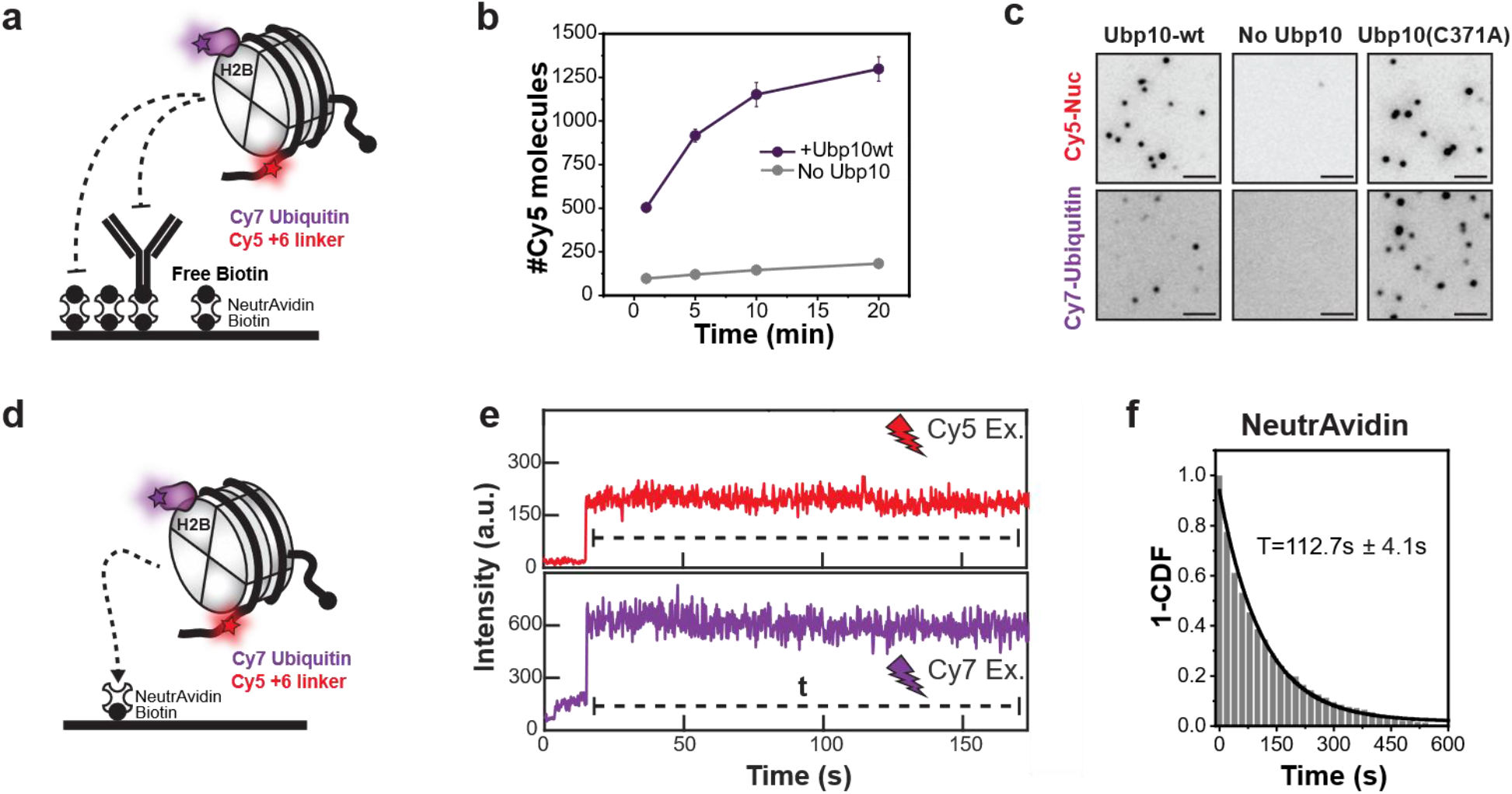
Controls for nucleosome binding specificity and Cy7 photobleaching kinetics. **a**, Schematic of the non-specific binding control assay. Slide surfaces were functionalized with NeutrAvidin and biotinylated anti-GFP antibody, followed by free biotin quenching to block unoccupied binding sites and prevent nucleosome adsorption. **b**, Surface-bound Cy5-nucleosome counts confirming Ubp10-mGold2t binding specificity over background. **c**, Representative images showing Cy5-nucleosome and Cy7-ubiquitin spots across different surface functionalizations. Wild-type Ubp10 displays reduced Cy7 colocalization relative to the catalytically dead mutant, whereas minimal background binding is observed in the absence of Ubp10 (scale bar, 5 µm). **d**, Schematic of real-time single-molecule flow experiments capturing biotinylated Ub-nucleosomes via direct NeutrAvidin pull-down. **e**, Representative single-molecule intensity trajectory showing persistent Cy5 and Cy7 signals upon nucleosome landing on NeutrAvidin. **f**, Survival probability plot (1-CDF) of Cy7 photobleaching lifetimes for nucleosomes directly immobilized on the TIRF surface in the absence of Ubp10.

**Extended Data Figure 3:**
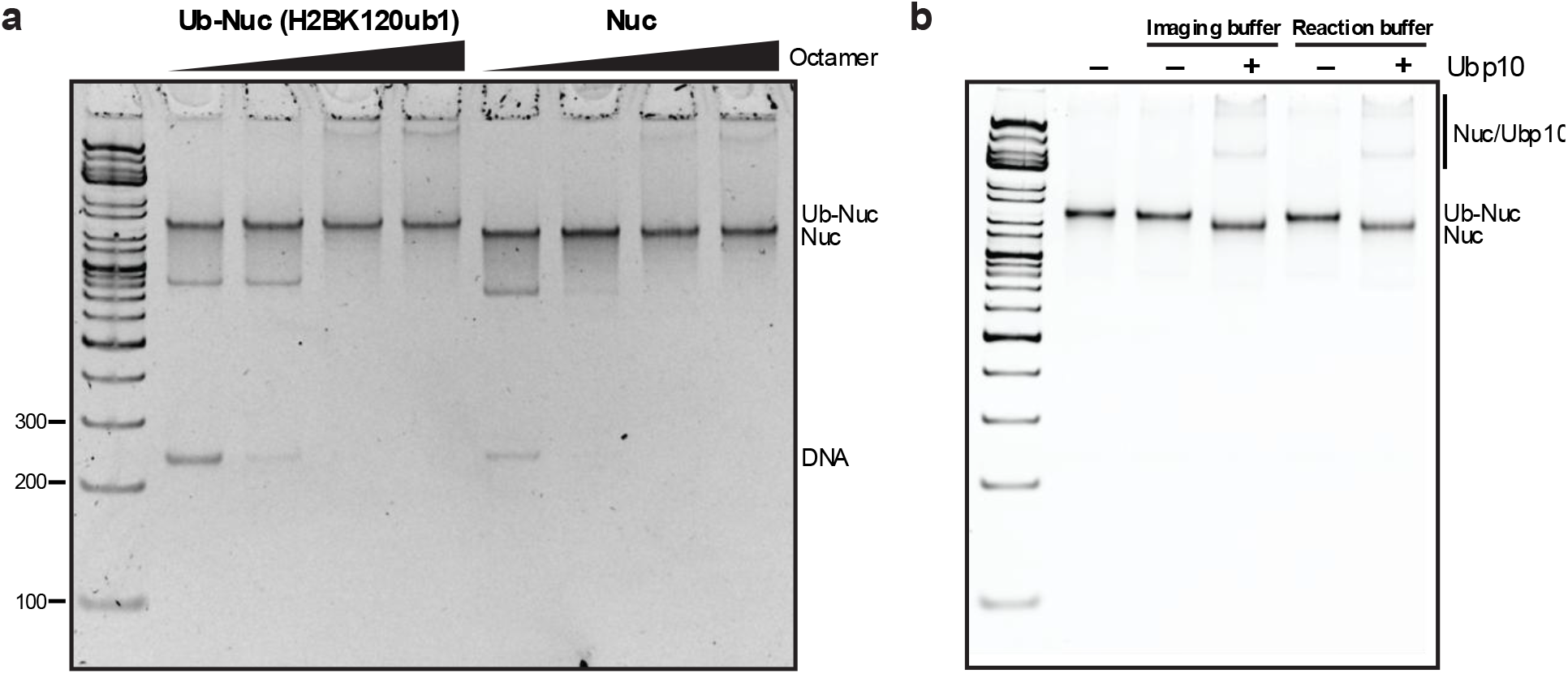
Large-scale reconstitution of ubiquitinated nucleosomes and bulk DUB activity assays. **a**, Native PAGE analysis of nucleosome reconstitution titrations using human H2BK120ub1 histone octamers (left) or unmodified human histone octamers (right) assembled onto a 233 bp DNA template containing a central Widom 601 sequence flanked by 43 bp linker arms. Ubiquitinated nucleosomes exhibit reduced electrophoretic mobility. **b**, Bulk deubiquitination assays performed in reaction buffer or imaging buffer (reaction buffer + Trolox) to validate Ubp10 activity at equimolar enzyme-to-substrate concentrations (333 nM each).

**Extended Data Figure 4:**
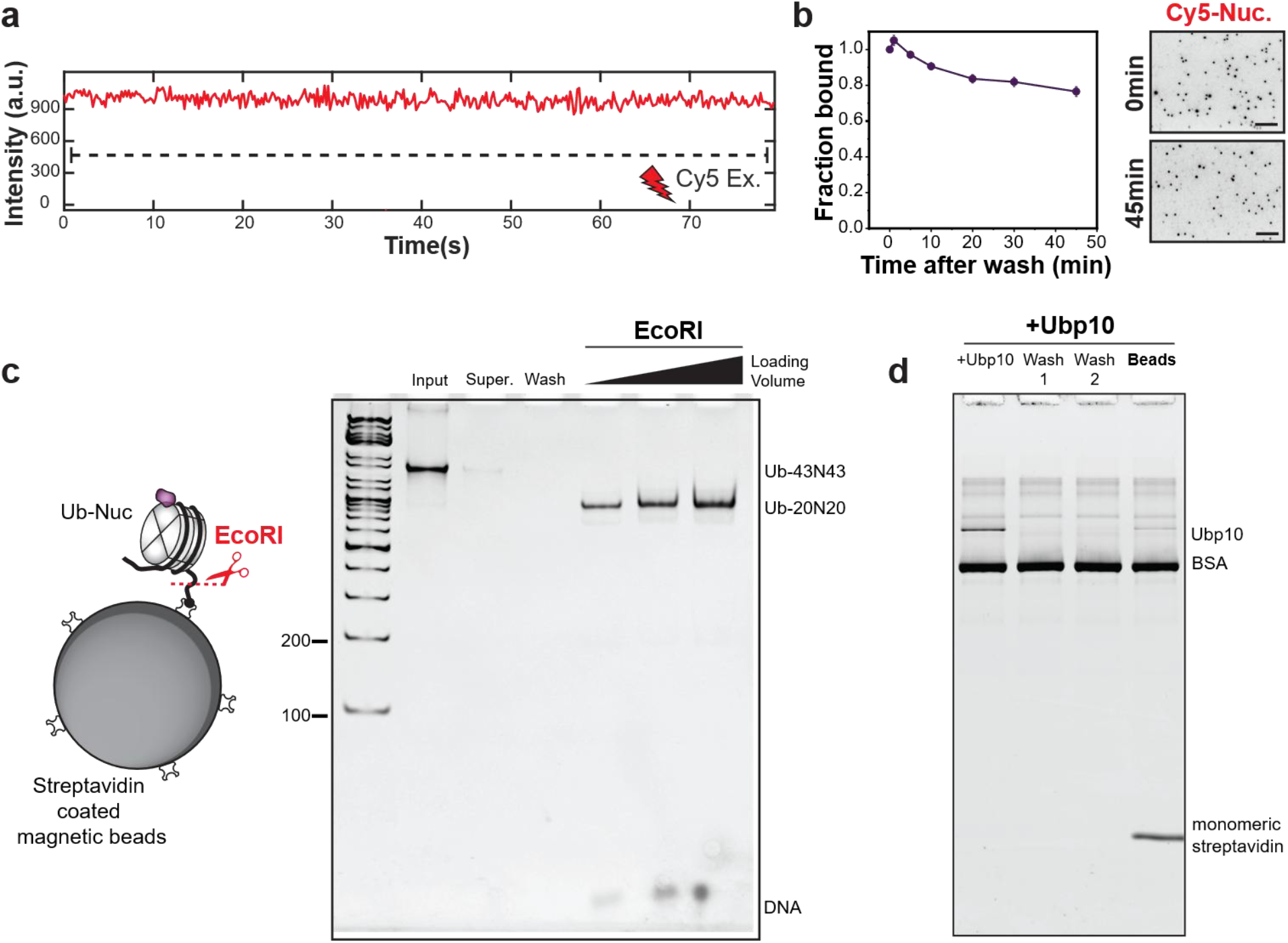
Stable engagement of nonubiquitinated nucleosomes to Ubp10 and magnetic bead controls. **a**, Representative Cy5 fluorescence intensity trajectory of long-lived, single nonubiquitinated nucleosome bound to surface-tethered Ubp10. **b**, Time-course tracking of Cy5-nonubiquitinated nucleosome stably bound over time (scale bar, 10 µm). **c**, Immobilization of Ub-nucleosomes (265 nM) on streptavidin magnetic beads. EcoRI restriction digestion within the linker DNA was performed to verify nucleosome integrity. **d**, Non-specific background control showing minimal Ubp10 adsorption to bare magnetic beads.

**Extended Data Figure 5:**
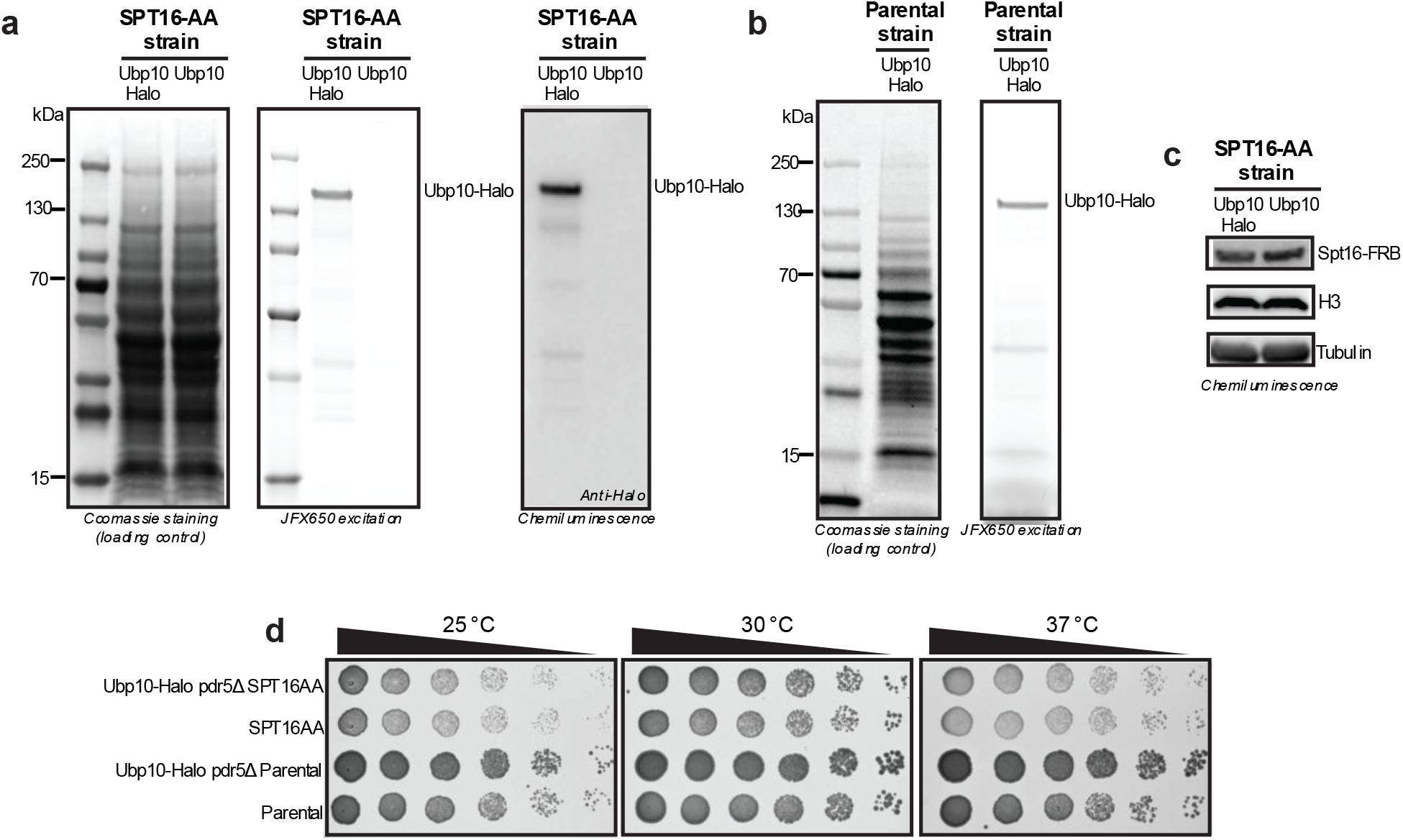
Validation of Ubp10-HaloTag expression, tag stability, and cell viability. **a**, SDS-PAGE and immunoblot analysis of Ubp10-HaloTag protein integrity. Cell lysates from Spt16-AA strains (+/-Ubp10-HaloTag) were prepared by trichloroacetic acid (TCA) precipitation following labeling with 7.5 nM JFX650-HaloTag ligand for 2 h at 30 °C. Samples were evaluated by total protein Coomassie staining (left), in-gel JFX650 fluorescence scanning (middle), and anti-HaloTag immunoblotting (right). **b**, Total protein Coomassie stain (left) and JFX650 fluorescence scan (right) of Ubp10-HaloTag parental control lysates lacking Spt16-FRB following dye treatment. **c**, Western blot analysis of Spt16, histone H3, and tubulin expression levels in untagged and Ubp10-HaloTag strains. **d**, Fivefold serial dilution growth assays comparing growth of Ubp10-HaloTag in Spt16-AA and parental control backgrounds on YPD agar at indicated temperatures for 2 days.

**Extended Data Figure 6:**
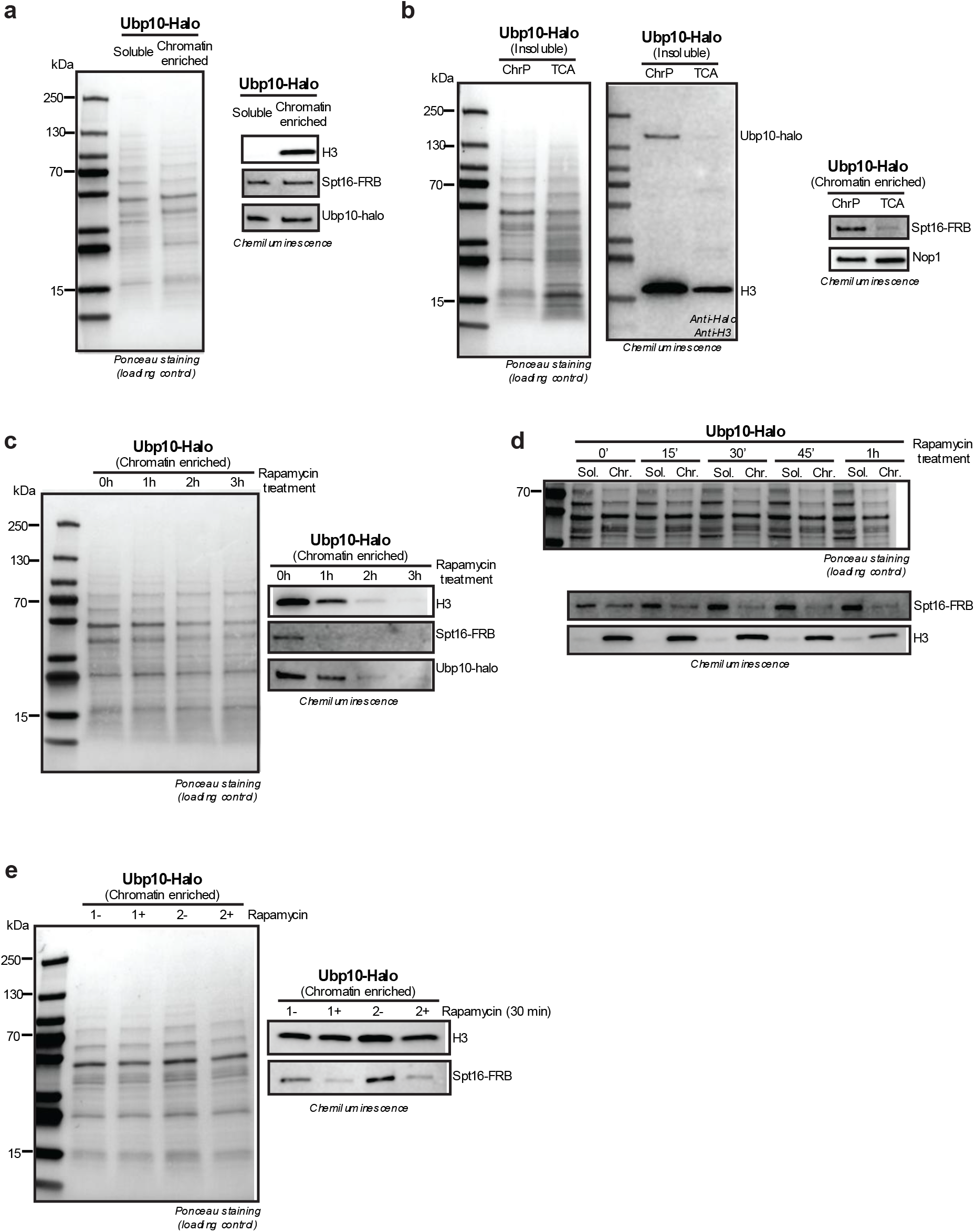
Optimization of rapamycin-mediated Spt16 depletion from chromatin. **a**, Chromatin fractionation separating free cytosolic/nuclear soluble proteins from chromatin/chromatin-bound species. Samples were evaluated by Ponceau staining (left), and immunoblotting (histone H3, Spt16 and HaloTag) (right). **b**, Protein recovery comparison of insoluble chromatin fractions. Insoluble chromatin pellets solubilized by MNase digestion were compared against total proteins recovered from the insoluble chromatin fraction by TCA precipitation. Signal levels of histone H3, Ubp10-HaloTag, Spt16, and Nop1 were evaluated by Ponceau staining (left) and immunoblotting (middle and right). Specific enrichment of Ubp10 and Spt16 in MNase-solubilized fractions validates that these recovered proteins represent true chromatin-bound species. **c**, Time-course evaluation of chromatin extractions at 0, 1, 2, and 3 h following 10 µg/mL rapamycin treatment. **d**, Higher-resolution time course (every 15 min for 1 h) of soluble and chromatin enriched fractions during rapamycin treatment. **e**, Chromatin extractions before and after 30 min Rapamycin treatment (n=2) confirming Spt16 depletion.

**Extended Data Figure 7:**
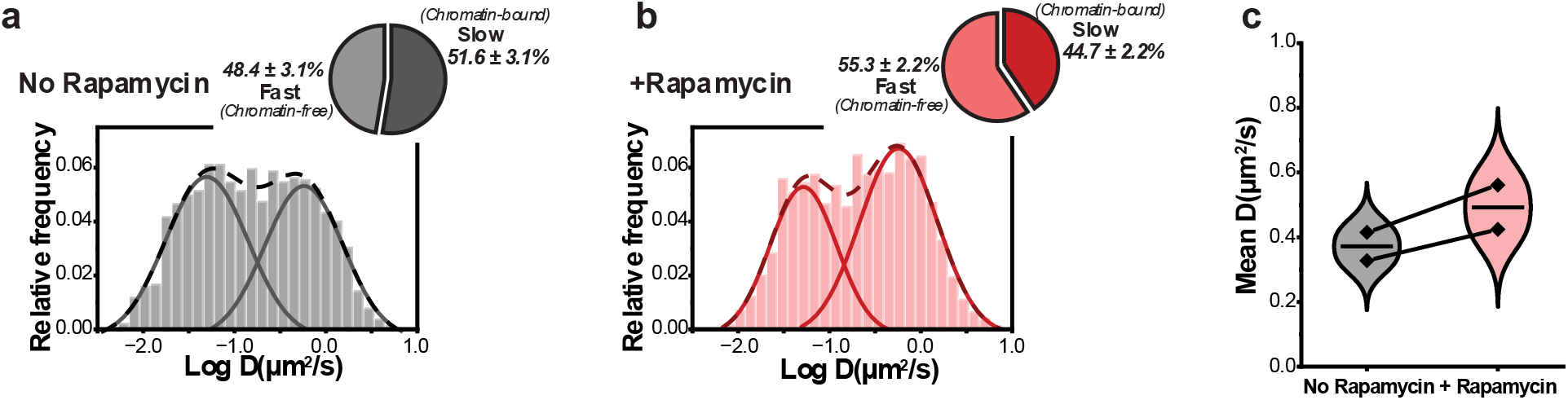
Ubp10 diffusion in the parental control strain upon rapamycin treatment. **a,b**, Two-state Gaussian fitting of log(D) diffusivity values for JF552-Ubp10 at (**a**) no rapamycin versus (**b**) + rapamycin (movies acquired 15–30 min after addition), showing relative proportions of bound and unbound fractions (dashed line: sum of Gaussians; solid lines: individual curves). **c**, Comparison of mean Ubp10 diffusion coefficients in the parental strain between conditions (n=2).

## Materials and Methods

### Protein expression and purification

Ubp10-WT and Ubp10(C371A) were expressed and purified using previously established methods^19^. Ubp10-mGold2t and Ubp10(C371A)-mGold2t, carrying a variation of the mYFP tag^41^ on the C-terminus separated by 3 repeats of glycine-serine (GS), were expressed and purified by Tierra Biosciences. During their production process an additional cleavable tag is added on the C-terminus. Consequently, the final mGold2t tagged Ubp10 proteins contain 7 extra amino acids. All proteins were aliquoted, stored at -80 °C and used only once to avoid freeze-thaw cycles.

Ubiquitin was expressed, purified and conjugated to Cy7 as previously described^20^. Nhp6a and Spt16/Pob3 heterodimers were expressed and purified using established protocols^57, 58^.

### Octamer refolding

Recombinant yeast histones were expressed and purified using previously established protocols^59^. For bulk DUB assays, recombinant human histone octamer with a native cleavable monoubiquitinated histone H2B (H2BK120ub1) was purchased from EpiCypher^60^ as a custom preparation.

### DNA construct formation

Two fluorescently labeled nucleosomal DNA constructs of the same length (233 bp) were generated. Each substrate carried the 601-c2 positioning sequence^61, 62^ and 43 bp of linker DNA at both ends. Following our previously established labeling scheme, the first biotinylated two-color DNA construct, used for smTIRF, contained Sulfo-Cy5 conjugated to the linker DNA (+6 bp) and Sulfo-Cy3 incorporated at superhelical location +1 (SHL)^20^. The second biotinylated DNA substrate, used for bulk assays, was only labeled with Sulfo-Cy5 at the SHL -7.

Amino-modified primers were labeled with their respective NHS-ester dyes (Lumiprobe). Free dye was removed using ethanol precipitation and the labeled primers were used for large scale PCR reactions. The two-color nucleosomal DNA construct was prepared as described previously^20^. The Sulfo-Cy5 labeled DNA was purified over a 6% native polyacrylamide (PAGE) column in a miniprep cell (Bio-Rad) using 0.5X TBE. Fractions were identified via agarose gel electrophoresis, collected and concentrated for storage at -20 °C.

### Nucleosome reconstitution

Standard salt gradient dialysis was used to reconstitute nucleosomes^59^. Yeast histone octamer was used for the two-color biotinylated DNA and ubiquitinated human histone octamer for the sulfo-Cy5 labeled DNA. Briefly, the desired amount of histone octamer was mixed with nucleosomal DNA in 10 mM Tris (pH 7.5), 2 M NaCl, 1 mM EDTA, and 1 mM β-mercaptoethanol. BSA was added for the two-color biotinylated DNA but omitted for the Sulfo-Cy5 construct to avoid background BSA signal during DUB analysis by SDS– PAGE. Reactions were placed on a 7 kDa MWCO dialysis button (Slide-A-Lyzer MINI dialysis unit) and dialyzed from high-salt buffer (10 mM Tris at pH 7.5, 2 M NaCl, 1 mM EDTA, 3.5 mM BME, and 0.02% NP-40) into low-salt buffer (10 mM Tris at pH 7.5, 50 mM NaCl, 1 mM EDTA, 3.5 mM BME, and 0.02% NP-40) at 4 °C overnight. Quality of assembled nucleosomes was verified using 6% native PAGE and reconstitutions were concentrated using a 10 kDa MWCO centrifugal unit.

Yeast reconstituted nucleosomes were ubiquitinated enzymatically by incubating them with human E1 and yeast Rad6 using Cy7-labeled ubiquitin (His tagged) and further purified using Ni-NTA beads as described previously^20^.

### smTIRF Microscope instrumentation

A custom-built prism-based total internal reflection fluorescence (TIRF) microscope was used for single-molecule experiments on a Nikon Eclipse Ti microscope^63^. Three solid-state lasers were used to excite sulfo-Cy3 (543 nm, Shanghai Dream Lasers Technology), sulfo-Cy5 (640 nm, Coherent) and sulfo-Cy7 (750 nm, Shanghai Dream Lasers Technology). Emitted fluorescence was collected by a Nikon water immersion 60X/1.27 NA (numerical aperture) objective and laser signal was blocked using a custom notch filter. A set of dichroic mirrors was used to separate signal into two or three channels and images were captured on an EMCCD (electron-multiplying charge-coupled device, Andor iXon 897) camera.

The home-built smCamera 2.0 software was used to record movies. Custom IDL (Interactive Data Language) and MATLAB scripts were used for background subtraction, spot selection, fluorescent signal quantification for single-molecule trajectories, smFRET and colocalization experiments (https://sites.google.com/site/taekjiphalab/resources).

### Quartz slide preparation

Quartz slides and glass coverslips were prepared as described previously^64^ with 98% methoxy polyethylene glycol (PEG) and 2% biotin-PEG. Amino-silanization and PEGylation were performed twice for increased passivation. Slides were assembled into flow chambers using double-sided tape and sealed with epoxy. For each experiment, single-molecule channels were washed with 100 µl of T50 buffer (10 mM Tris-HCl pH 7.5, 50 mM NaCl) and later the surface was functionalized by flowing 40 µl of NeutrAvidin (0.2 mg/ml in T50 buffer) incubated for 1 min. Free NeutrAvidin was removed by washing with T50 buffer, and the channel was immediately used for single-molecule immobilization.

### Nucleosome immobilization conditions for smTIRF imaging

Fluorescently labeled 3-color ubiquitinated nucleosomes were diluted to ∼15 pM in Buffer 1 (25 mM HEPES-KOH pH 7.6, 0.37 mM EDTA, 5% Glycerol, 0.017% NP-40, 70 mM KCl, 3.6 mM MgCl2 and 100 μg/ml BSA) and incubated for 2 min on the NeutrAvidin coated chambers. Free nucleosomes were washed off with 100 µl of Buffer 1 and 100 µl of imaging buffer (Buffer 1 supplemented with an oxygen-scavenging system containing 0.8% w/v dextrose, 2 mM Trolox, 1 mg/ml glucose oxidase, and 500 U/ml catalase). Nucleosomes were imaged right after immobilization prior to the addition of additional proteins.

### Ubp10 immobilization conditions for smTIRF imaging

Biotin anti-GFP antibody (30 nM) was diluted into T50 buffer supplemented with 0.1 mg/ml BSA (T50-BSA) and flowed into NeutrAvidin functionalized chambers. After 10 min of incubation, free antibody was washed off with 100 µl of T50-BSA followed by 100 µl of reaction buffer (20 mM Tris-HCl pH 7.5, 100 mM NaCl, 1 mM EDTA pH 8.0, 10% Glycerol and 0.1 mg/ml BSA). Ubp10-mGold2t was diluted in reaction buffer to 4 nM, flowed into the channel, incubated for 10 min and later washed using reaction buffer. The NeutrAvidin on the single-molecule surface was neutralized with two consecutive 10 min incubations of 100 µM free biotin diluted in reaction buffer. Chambers were washed one last time with reaction buffer and used for nucleosome pulldown experiments.

### Three-color imaging for FRET

Alternating laser excitation (ALEX) was used to excite Cy3, Cy5 and Cy7 dyes. A total of 10 short movies (200 ms exposure time, 30 frames each) were acquired. Cy3-Cy5 colocalized spots were analyzed to calculate the FRET efficiency of single nucleosomes^20^. Additionally, Cy7 positive molecules were quantified to determine the fraction of ubiquitinated nucleosomes under each condition.

### Two-color imaging for colocalization

Ten short movies were taken with a total of 20 frames each (200 ms exposure), the first 10 for Cy5 excitation followed by 10 for Cy7 excitation. The Cy7 intensity histograms on Cy5 selected spots were plotted to evaluate nucleosome ubiquitination state and quantify DUB activity.

During real-time flow experiments, long movies of 5000 frames were taken using Cy5 and Cy7 ALEX (200 ms exposure). Nucleosome binding was counted as the concurrent appearance of Cy5 and Cy7 signal throughout the imaging time. The Cy7 dwell time was collected by manual inspection of single-molecule trajectories using custom MATLAB scripts and the lifetime was extracted by fitting 1-CDF (cumulative distribution function) of the dwell time to a single or double exponential decay function.

### Steady-state measurements of DUB activity on immobilized nucleosomes

Single-molecule Ubp10 DUB activity was examined by immobilizing nucleosomes in the flow chamber and incubating them in Imaging Reaction Buffer (reaction buffer supplemented with the oxygen-scavenging system containing 0.8% w/v dextrose, 2 mM Trolox, 1 mg/ml glucose oxidase, and 500 U/ml catalase) with increasing concentrations of Ubp10 only (10, 50, 100, 200 pM), FACT only (1 µM Spt16/Pob3, 2 µM Nhp6a), or Ubp10 (10 pM) + FACT (1 µM Spt16/Pob3, 2 µM Nhp6a). Short movies were taken at different timepoints using three-color imaging setup. Cy7 colocalized signal to Cy3-Cy5 positive spots was used to calculate the fraction of ubiquitinated nucleosomes. The progress of the DUB reaction was monitored by dividing the fraction of Ub-Nucs at a specific time by the fraction of Ub-Nucs before exposure to Ubp10. The Cy3 and Cy5 fluorescence intensity was used to determine FRET efficiency before and after Ubp10 incubation. Similar experiments were performed by injecting 250 nM or 1 µM of the catalytically inactive Ubp10(C371A).

### Ub-Nucleosome pulldown on surface-tethered Ubp10

Three-color Ub-nucleosomes were diluted in imaging buffer and flowed to the Ubp10-immobilized chambers. The number of nucleosomes binding to Ubp10 was counted as the number of Cy5 spots engaging the TIRF surface in a total of 10 short movies per timepoint. Non-specific binding was assessed by incubating Ub-nucleosomes to chambers prepared similarly but omitting the Ubp10-mGold2t immobilization step.

### Ub-Nucleosome real-time flow and dwell time measurement

For real-time measurements, 50 µl of Ub-Nuc diluted in imaging buffer was injected at the start of recording (0 s) using a syringe pump at 1 ml/min flow rate. As a separate experiment, the Cy7 photobleaching lifetime was similarly followed by flowing and recording Ub-Nuc binding to NeutrAvidin-only functionalized chambers.

### DUB bulk assays and EMSA conditions

Cy5-labeled reconstituted Ub-nucleosomes (333 nM) were mixed with 333 nM Ubp10 (His6-tag cleaved) in Reaction Buffer at RT using Protein LoBind Tubes (Eppendorf). Time-courses were recorded by removing 3 µl of the reaction mix at the indicated time points and quenching with LDS buffer (Bio-Rad). Samples were heated for 5 min at 95 °C and loaded onto 16% SDS-PAGE gels (Invitrogen, Novex Tris-Glycine), stained with Flamingo Fluorescent Protein Gel Stain (Bio-Rad) and imaged using a Typhoon scanner.

To determine if nucleosomes remained stable in DUB bulk reaction conditions, 2 µl of the leftover reaction mix (after 1 hour of incubation with Ubp10) was run on a 6% native PAGE. Two additional controls were included, nucleosomes without Ubp10 diluted in Reaction Buffer and incubated for an hour at RT, and nucleosomes kept on ice. Gel was run at 110 V for 1 h using 0.5X TBE and stained with EtBr.

### Magnetic bead nucleosome immobilization and DUB assay

Dynabeads MyOne Streptavidin T1 (Invitrogen) were prepared as follows. First, 40 µl of resuspended beads were transferred to a 1.5 ml Eppendorf tube and placed on a magnet. Beads were washed three times with 200 µl of Washing Buffer (5 mM Tris-HCl pH 7.5, 0.5 mM EDTA, 1 M NaCl), followed by three 200 µl washes using Bead Reaction Buffer (Reaction Buffer supplemented with 0.017% NP-40 and 100 μg/ml BSA, included to minimize nonspecific adsorption of Ubp10 to the beads). All subsequent wash, protein and nucleosome dilution was performed using Bead Reaction Buffer. Beads were resuspended in 25 µl of 265 nM Cy5 labeled Ub-Nuc and incubated with shaking for 5 min. Beads were placed on a magnet and 5 µl of supernatant was collected. Next, beads were washed three times with 50 µl of Bead Reaction buffer and resuspended in 25 µl of 265 nM Ubp10 (2 min, with shaking). Beads were placed on a magnet and 5 µl of the reaction was saved to run on a gel before discarding the rest. Beads were washed twice with 25 µl, left for 20 minutes at RT and placed on a magnet to take 10 µl. Lastly, every sample collected and the final beads were denatured with LDS buffer and heated prior to loading onto 16% SDS-PAGE gels.

A control reaction was conducted to determine the extent of non-specific Ubp10 binding to the magnetic beads, in which the nucleosome immobilization step was omitted and replaced with Reaction Buffer lacking biotinylated nucleosomes. Additionally, to ensure that nucleosomes were stable under the on-bead DUB assay conditions we digested the linker DNA of bead-immobilized nucleosomes using EcoRI. Briefly, after incubating nucleosomes with beads and three rounds of wash, they were resuspended with 1X CutSmart buffer and 1 µl of EcoRI (10 min incubation at 37 °C). Beads were placed on magnet, and the supernatant was run on a 6% native PAGE gel.

### Off-rate nucleosome-Ubp10 measurements

Nucleosomes that were pulled down by surface-tethered Ubp10 were washed by flowing 150 µl of imaging buffer with or without FACT (1 µM Spt16/Pob3, 2 µM Nhp6a) onto the single-molecule chamber. The number of Cy5-labeled nucleosomes remaining on the surface was then tracked over time by acquiring short movies under red (640 nm) excitation at increasing time points after the wash. For each time point, Cy5 spots were counted across 10 movies taken at different positions in the channel, and the fraction of nucleosomes remaining bound was calculated by normalizing to the number of Cy5 spots counted immediately before the wash.

### Yeast strains

The *Saccharomyces cerevisiae* background strains used in this study were kindly provided by Kevin Struhl. The Spt16 Anchor-Away strain (yFR1478) was generated in François Robert’s laboratory. The parental HHY168 control strain was obtained from Euroscarf (Supplementary Table S2). Standard yeast transformation protocols were used to delete the Pdr5 ABC transporter gene (to ensure intracellular retention of HaloTag ligands) and to fuse the HaloTag sequence to the C-terminus of Ubp10 via a (SG_4_)_2_ flexible linker.

For HaloTag integration, the plasmid pTSK561 was used (a gift from Tatiana Karpova & Gunjan Mehta; Addgene plasmid #190816). The Pdr5 locus was disrupted via homologous recombination using a hygromycin resistance cassette (hphMX6) amplified from pFA6a-mCherry-hphMX6 (a gift from Jian-Qiu Wu; Addgene plasmid #105156). List of primers is provided in Supplementary Table S3.

### Yeast Growth and Spot Assays

To assess whether C-terminal HaloTag fusion altered cell viability, serial dilution spot assays were performed comparing Ubp10-HaloTag strains to their respective parental controls. Saturated overnight cultures grown in YPD were adjusted to equivalent densities, 5-fold serially diluted and spotted onto YPD agar plates for incubation at 25 °C, 30 °C, and 37 °C. Plates were photographed after 48 h of growth.

### Protein Extraction and HaloTag Integrity Analysis

Yeast cultures grown to early log phase (OD_600_ = 0.3-0.4) were incubated with 7.5 nM JFX650 HaloTag ligand for 2 h at 30 °C. Total protein was extracted using trichloroacetic acid (TCA) precipitation^65^. Protein concentrations were determined using the detergent-compatible DC Protein Assay Kit II (Bio-Rad).

For each sample, 100 µg of total protein was resolved on a 4–20% SDS-PAGE gel. One gel was imaged for in-gel JFX650 fluorescence on an Amersham Typhoon scanner (Cy5 excitation channel) and then stained with Coomassie as a loading control. Proteins from a second, identically loaded gel were transferred to nitrocellulose membranes, stained with Ponceau S to confirm transfer, and blocked prior to antibody incubation. Primary antibodies used were: anti-Spt16 (Cusabio, CSB-PA022955XA01SVG; 1:2,000), anti-histone H3 (Abcam, ab1791; 1:4,000), anti-HaloTag (Promega, G9211; 1:1,000), and anti-alpha-tubulin (Abcam, ab184970; 1:5,000). Horseradish peroxidase (HRP)-conjugated anti-mouse (1:2,000) and anti-rabbit (1:4,000) secondary antibodies were applied, and chemiluminescent signals were captured on an Azure Biosystems imager.

### Native Chromatin Fractionation

Yeast cultures were grown to early log phase (OD_600_ = 0.3 - 0.4) and harvested before or after rapamycin treatment (10 µg/mL) across indicated time courses. For each condition, 100 mL of cell culture was collected. Native chromatin extraction was performed as previously described^66^ with minor modifications. Briefly, cell pellets were washed twice with 1 mL Nucleosome Isolation Buffer (NIB: 20 mM HEPES pH 7.5, 250 mM sucrose, 10 mM MgCl_2_, 75 mM NaCl, supplemented with protease and phosphatase inhibitor cocktails). Pellets were resuspended in an equal volume of NIB buffer and glass beads, then lysed by five cycles of 45 s vortexing alternating with 3 min on ice.

Lysates were separated from glass beads and clarified by centrifugation to yield the soluble fraction. The insoluble chromatin pellet was washed twice with 1 mL NIB buffer (1,000 x g for 4 min at 4 °C), resuspended in 200 µL NIB buffer containing 5 mM CaCl_2_, and digested with micrococcal nuclease (MNase, 180 units) for 10 min at 37 °C. Digestion was quenched with EDTA (10 mM final concentration), and NaCl was added to a final concentration of 150 mM. Solubilized chromatin was recovered by centrifugation at 16,000 x g for 10 min at 4 °C, and protein concentrations were quantified using the DC Protein Assay Kit II (Bio-Rad). Equal amounts of protein (10 µg) from soluble and chromatin-enriched fractions were resolved on 4–20% SDS-PAGE gels, transferred to nitrocellulose membranes, Ponceau S-stained, and probed with anti-H3, anti-Spt16, anti-HaloTag, and anti-Nop1 (ThermoFisher Scientific, 28F2; 1:1,000) primary antibodies.

### Live-Cell Single-Particle Tracking (spTracking) Sample Preparation

Yeast cultures were grown to early log phase (OD_600_ = 0.2) in Complete Synthetic Medium (CSM) and labeled with 50 nM JF552-HaloTag ligand for 2 h at 30 °C. Saturating dye conditions allowed distinct spatial separation of the dense nucleolar Ubp10 pool from the nucleoplasmic population. Cells were harvested by centrifugation (1,000 x g for 2 min), washed four times, and resuspended in fresh CSM.

Resuspended cells were loaded into a microfluidic flow chamber assembled by mounting a self-adhesive sticky-Slide VI^0.4^ (ibidi, 80608) onto a concanavalin A-coated coverslip (Fisherbrand, 12-545-M). Unattached cells were removed by washing with CSM. Cells were incubated with 100 µg/ml Hoechst for 5 min and washed with CSM. To induce Spt16 Anchor-Away depletion during live imaging, CSM containing 10 µg/mL rapamycin was flowed directly to the chamber.

### Widefield single-particle tracking

For camera-based single-particle tracking of Ubp10-HaloTag fluorescently tagged with a JF552-Halotag ligand, we utilized a Nikon Eclipse microscope equipped with Nikon perfect focus system, with a solid-state white light source (Lumencor, SOLA SE II 365) and a custom total internal reflection fluorescence microscopy (TIRFM) module. The microscope was controlled by the Nikon NIS-Elements software. Highly Inclined and Laminated Optical sheet (HILO) was configured using a 100x oil immersion objective (Olympus 1.40 Oil UPlanSApo) with a custom dichroic (Chroma, zt 405/488/561/647/752rpc-UF2) and a 561 nm laser (MPB Communications, F-04306-102). To image the Hoechst channel, the white light source together with a Nikon filter cube (Nikon 96359 DAPI) was utilized.

First, widefield images of the Hoechst labeled sample and the densely labeled sample JF552 channel were acquired^49^. Then, JF552 was driven into a dark state by laser excitation. Because JF552 stochastically reverts to a fluorescent state, this produces super-resolution-like blinking and enables sparse single-molecule tracking. SpTracking images were acquired every 10 ms using a sCMOS camera (ORCA Flash 4.0, Hamamatsu Photonics) with a 2x2 binning mode at 300 mW laser power for around 2 min.

### SpTracking analysis

The spTracking movie files were localized and the diffusion parameters analyzed using the GEMspa plugin in napari^67^. For the diffusion analysis a minimum of 7 data points per trace was required and an apparent diffusion coefficient (D_app_) was fitted to the data within the GEMspa plugin:

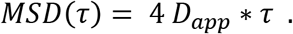

To extract the diffusion coefficients of Ubp10 interacting with chromatin and exclude nucleolar Ubp10, we manually annotated masks according to the Hoechst channel and excluded areas exhibiting a strong (nucleolar) signal on the JF552 channel. Also, cells exhibiting a very strong JF552 signal, due to e.g. stress, were excluded.

For the comparison of Ubp10 diffusion before and after nuclear Spt16 depletion, movies were assigned to two conditions. The “no rapamycin” condition comprised all movies acquired from a sample before rapamycin-containing medium was flowed into the chamber. The “+ rapamycin” condition comprised movies acquired between 15 and 30 min after rapamycin addition, pooled per sample; movies recorded during the first 15 min of treatment were not included because Spt16 depletion from chromatin is still incomplete at these early times (Extended Data Fig. 6d). Trajectories from all movies within a condition were combined to build the log(D_app_) distributions (Fig. 4i,j and Extended Data Fig. 7a,b), and the mean apparent diffusion coefficient of each condition (Fig. 4h and Extended Data Fig. 7c) was calculated from the same pooled trajectories. Each independent replicate consisted of one sample imaged before and after rapamycin addition, so that the no-rapamycin and + rapamycin values were compared in a paired manner.

To estimate the error on occupancy, we first fit a sum of two Gaussians to the pooled log10(D) distribution across all stages to obtain fixed diffusion-coefficient peak positions and widths. Holding these fixed, we re-fit the two Gaussian amplitudes 1,000 times on randomly subsampled fractions of 25% and took the standard deviation of the resulting occupancy estimates as the bootstrap error.

### Statistics and reproducibility

All single-molecule TIRF experiments were performed in three independent replicates. The fraction of nucleosomes bound, and the fraction of ubiquitinated nucleosomes were normalized to the initial value measured at time zero in each replicate and are reported as the mean ± standard error of the mean (SEM) across replicates. For dwell-time measurements, the duration of the fluorescent signal was extracted from individual single-molecule trajectories pooled from three replicates, with at least 400 events analyzed per condition, and lifetimes were obtained by fitting the 1-CDF of the pooled dwell times as described above. Live-cell single-particle tracking experiments were performed in five independent replicates for Spt16 Anchor Away strain and in duplicate for parental control, and bootstrap errors on state occupancies were estimated as described above. Statistical significance between conditions in single-molecule TIRF and single-particle tracking experiments was assessed using a two-tailed Student’s t-test: unpaired for comparisons between conditions in single-molecule TIRF experiments and paired on the per-replicate values before and after rapamycin in spTracking experiments. Bulk deubiquitination assays and magnetic-bead pulldown assays were performed in triplicate, and representative gels are shown.

## Acknowledgements

We thank Xinyu A. Feng for advice at the outset of this project. We thank Matthew F. Poyton for his contribution to the design of the three-color ubiquitinated nucleosome construct used in our previous work and in this study, and for preliminary observation of Ubp10 activities in those nucleosomes. We thank Tim Formosa, Zaily Connell, and Laura McCullough for preparing purified FACT for the single-molecule experiments. We are grateful to Kevin Struhl for hosting R.M.-U. in his laboratory for the yeast *in vivo* work, and to Catherine Maddox of the Struhl laboratory for preparing reagents. We thank Carl Wu and Yick Hin Ling for sharing yeast strains used in the initial proof-of-concept single-particle tracking experiments in the Ha laboratory and for advice on yeast genetics, and Stephen Buratowski and Yoo Jin Joo for technical advice on yeast biochemistry. We thank Stephen D. Fried for discussion and suggestions. J.Z. was supported by a Feodor Lynen Research Fellowship from the Alexander von Humboldt Foundation. This work was supported by the National Institute of General Medical Sciences (GM130393 to C.W. and GM122569 to T.H.). T.H. is an investigator of the Howard Hughes Medical Institute.

## Author contributions

R.M.-U., T.H. and M.N. designed the experiments. R.M.-U. performed single-molecule TIRF experiments, bulk biochemical assays, and in vivo yeast work. J.Z. performed live-cell single-particle tracking experiments and analysis. M.N. purified Ubp10 proteins. C.W. supervised work carried out by M.N. R.M.-U. and J.Z. wrote software for data analysis. R.M.-U. and T.H. wrote the manuscript with input from all authors. C.W., M.N., and J.Z. edited the manuscript. T.H. supervised the project.

## Data availability

Source data for all figures, including unprocessed gel and blot images (Fig. 3c and Extended Data Figs. 3, 4c,d, 5 and 6) and the numerical values underlying all graphs, are provided with this paper. Raw single-molecule TIRF movies, extracted single-molecule trajectories and live-cell single-particle-tracking datasets will be deposited in Zenodo upon publication. Yeast strains, plasmids and DNA constructs generated in this study are available from the corresponding author upon request. Plasmids pTSK561 (Addgene 190816) and pFA6a-mCherry-hphMX6 (Addgene 105156) are available from Addgene.

## Code availability

Custom MATLAB, IDL and Python scripts used for single-molecule TIRF analysis (spot selection, trajectory extraction, smFRET, dwell time and colocalization) and for single-particle-tracking analysis (trajectory filtering and bootstrap estimation of state occupancies), together with the parameter settings used in GEMspa, are provided as Supplementary Software with this submission and will also be publicly shared upon publication.

## Supplementary Tables

**Supplementary Table S1.**
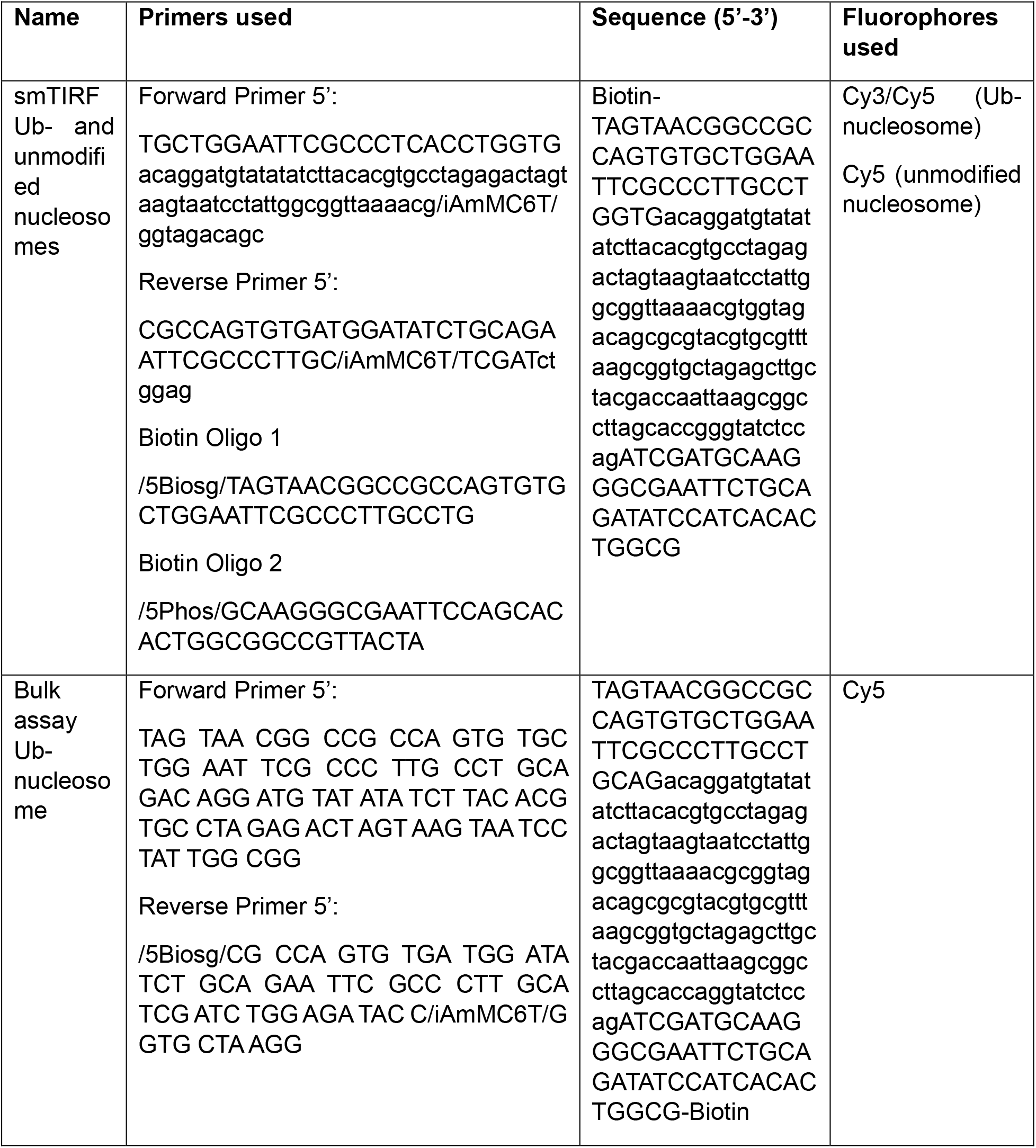
DNA and primers used for Ub- and unmodified nucleosomes (smTIRF and bulk assays)

**Supplementary Table S2.**
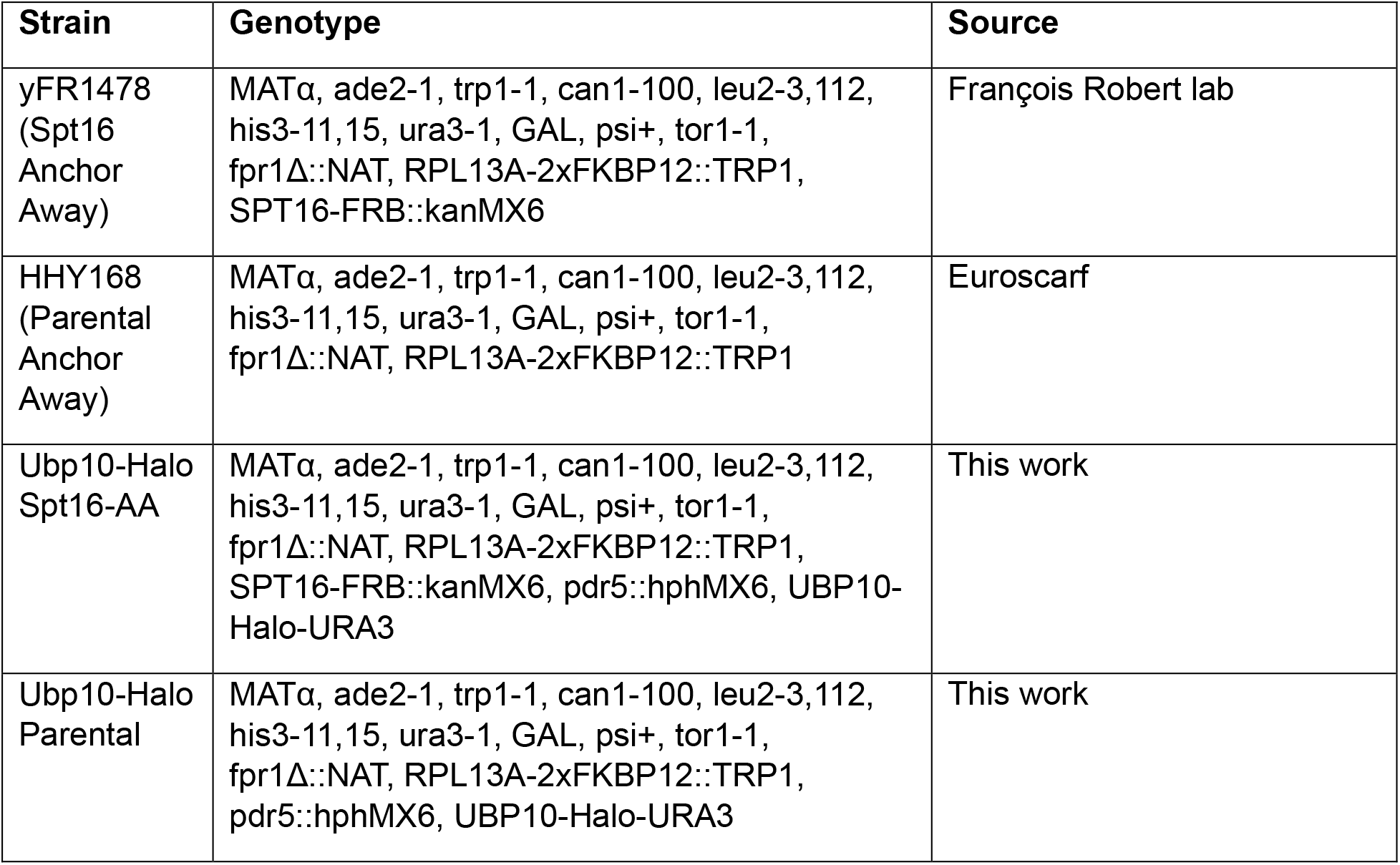
List of yeast strains used in this study.

| **Strain** | **Genotype** | **Source** |
|---|---|---|
| yFR1478 (Spt16 Anchor Away) | MATα, ade2-1, trp1-1, can1-100, leu2-3,112, his3-11,15, ura3-1, GAL, psi+, tor1-1, fpr1Δ::NAT, RPL13A-2xFKBP12::TRP1, SPT16-FRB::kanMX6 | François Robert lab |
| HHY168 (Parental Anchor Away) | MATα, ade2-1, trp1-1, can1-100, leu2-3,112, his3-11,15, ura3-1, GAL, psi+, tor1-1, fpr1Δ::NAT, RPL13A-2xFKBP12::TRP1 | Euroscarf |
| Ubp10-Halo Spt16-AA | MATα, ade2-1, trp1-1, can1-100, leu2-3,112, his3-11,15, ura3-1, GAL, psi+, tor1-1, fpr1Δ::NAT, RPL13A-2xFKBP12::TRP1, SPT16-FRB::kanMX6, pdr5::hphMX6, UBP10-Halo-URA3 | This work |
| Ubp10-Halo Parental | MATα, ade2-1, trp1-1, can1-100, leu2-3,112, his3-11,15, ura3-1, GAL, psi+, tor1-1, fpr1Δ::NAT, RPL13A-2xFKBP12::TRP1, pdr5::hphMX6, UBP10-Halo-URA3 | This work |

**Supplementary Table S3.**
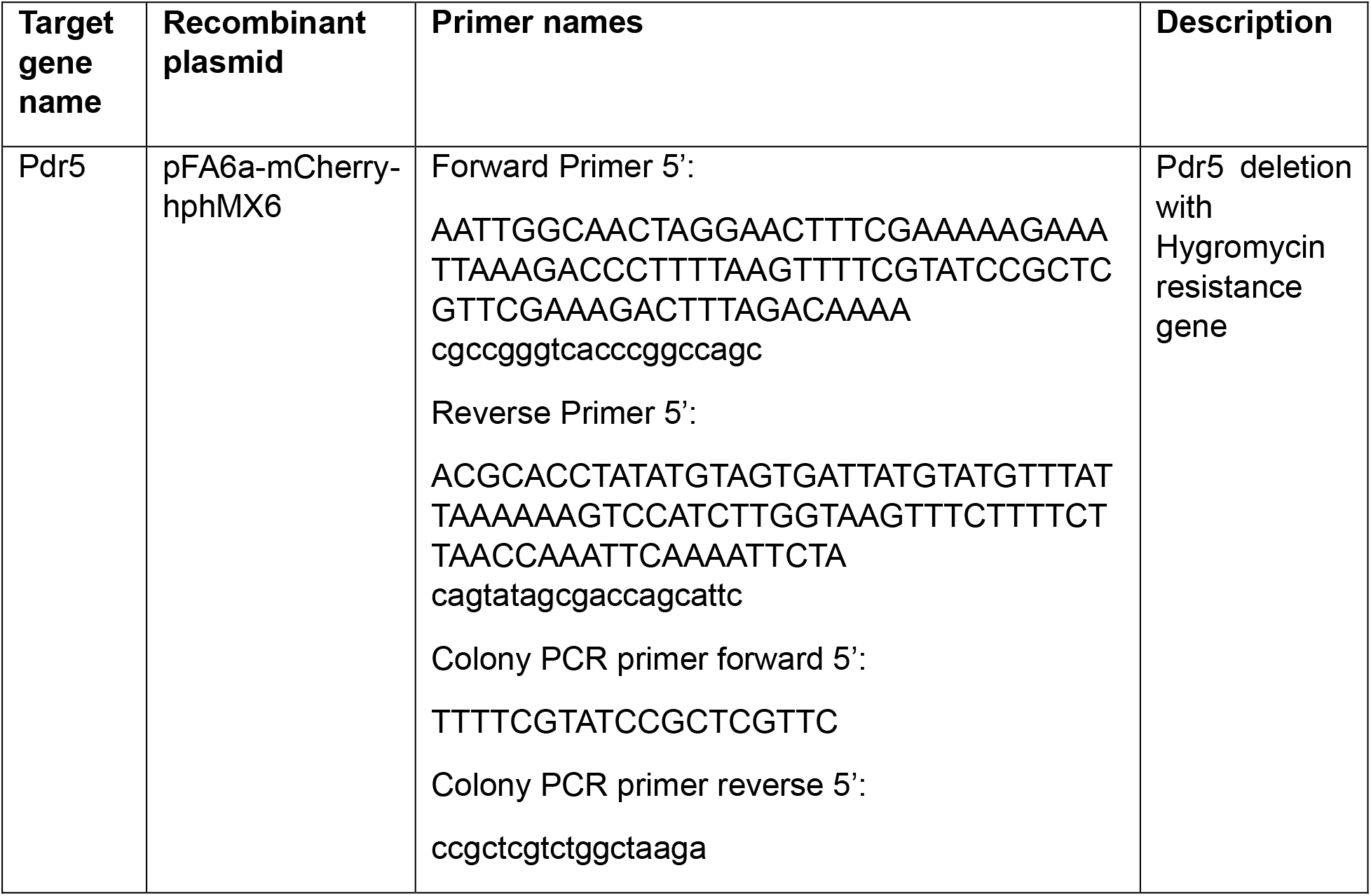
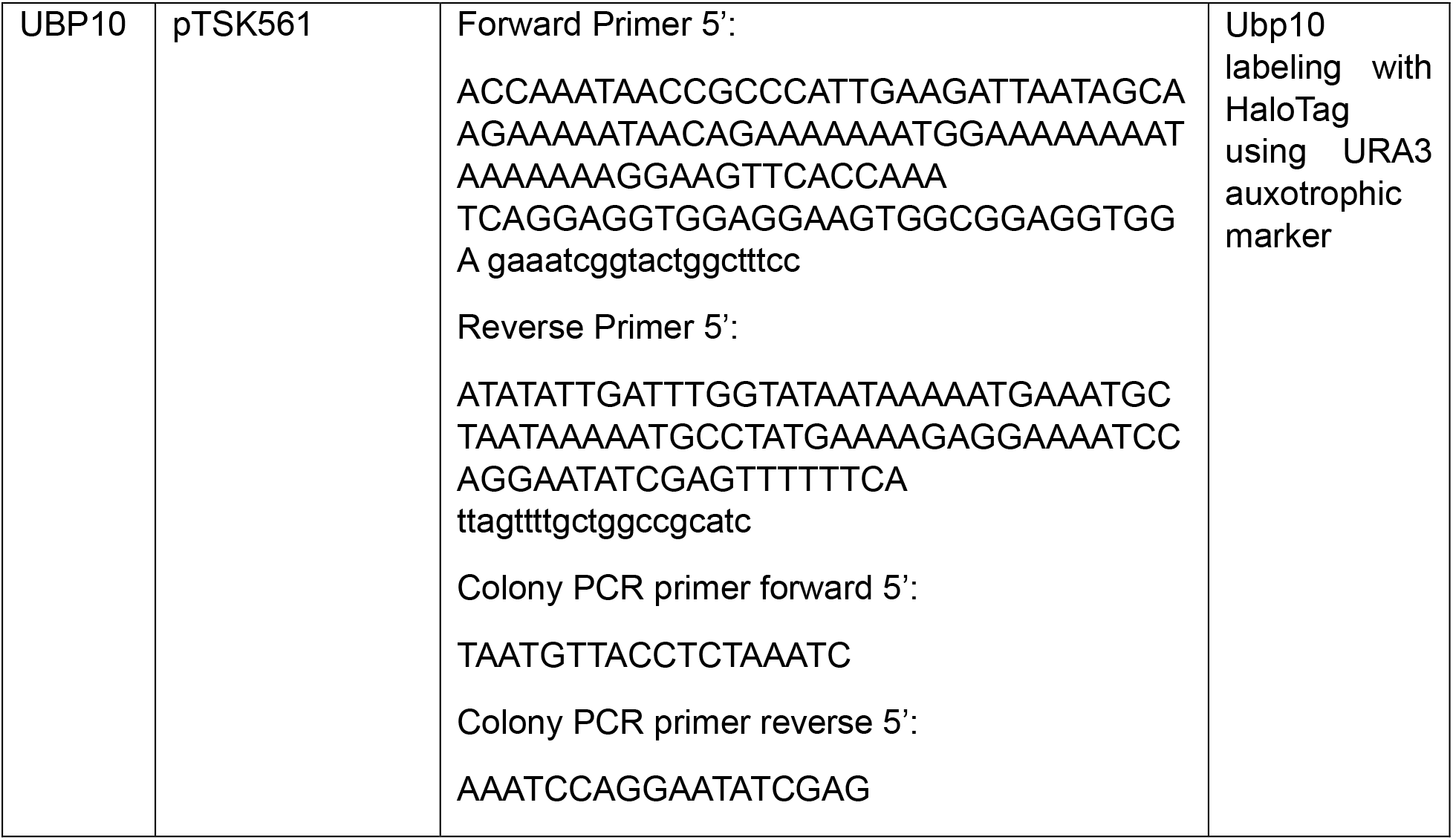
Oligonucleotides used for yeast strain construction.

### Materials

| Reagent or resource | Source | Identifier |
|---|---|---|
| Antibodies |  |  |
| SPT16 Antibody | Cusabio | CSB-PA022955XA01SVG |
| NOP1 Monoclonal Antibody (28F2) | ThermoFisher Scientific | MA1-10025 |
| Anti-Histone H3 antibody | Abcam | ab1791 |
| Anti-HaloTag® Monoclonal Antibody | Promega | G9211 |
| Anti-GFP (rabbit) Antibody Biotin Conjugated | Rockland | 600-406-215 |
| Recombinant DNA |  |  |
| pFA6a-mCherry-hphMX6 | Addgene | 105156 |
| pTSK561 | Addgene | 190816 |
| Chemicals, peptides and recombinant proteins |  |  |
| Concanavalin A from Canavalia ensiformis (Jack bean) | Sigma-Aldrich | L7647 |
| Rapamycin | MCE MedChemExpress | HY-10219 |
| Janelia Fluor® 552 in vivo HaloTag® Ligand Kit | Promega | CS3495C04 |
| 4–20% Mini-PROTEAN® TGX™ Precast Gel | Bio-Rad | 4561093 |
| Micrococcal Nuclease | New England Biolabs (NEB) | M0247S |
| Ponceau S Staining Solution | ThermoFisher Scientific | A40000278 |
| Q5<sup>®</sup> High-Fidelity DNA Polymerase | New England Biolabs (NEB) | M0491L |
| Phusion High-Fidelity PCR Master Mix with HF Buffer | New England Biolabs (NEB) | M0531L |
| Recombinant Human, H2BK120ub1 octamer | EpiCypher | N/A |
| EcoRI-HF | New England Biolabs (NEB) | Cat# R3101 |
| Thermo Scientific™ Slide-A-Lyzer™ MINI Dialysis Device, 7K MWCO | Fisher Scientific™ | PI69560 |
| Flamingo™ Fluorescent Protein Gel Stain | Bio-Rad | 1610491 |
| UltraPure™ BSA (50 mg/mL) | ThermoFisher Scientific | AM2618 |
| Dynabeads™ Streptavidin Magnetic Beads | Invitrogen | 65601 |
| NeutrAvidin Protein | Thermo Fisher Scientific | 31000 |
| Catalase from bovine liver | Sigma-Aldrich | C3155 |
| Glucose Oxidase from Aspergillus niger | Sigma-Aldrich | G2133 |
| Quartz slides | G Finkenbeiner | 100 |
| Micro Cover Glasses | VWR | 48393 |
| N-(2-Aminoethyl)-3-aminopropyltrimethoxysilane | EMD Millipore | 8.19172.0100 |
| Alconox Powder Detergent | Sigma-Aldrich | 242985 |
| Trolox ®, 97% | Fisher scientific | AC218940050 |
| Sulfo-Cyanine5 NHS ester | Lumiprobe | 23320 |
| Sulfo-Cyanine3 NHS ester | Lumiprobe | 21320 |
| Sulfo-Cyanine7 NHS ester | Lumiprobe | 15320 |
| Potassium hydroxide | Sigma-Aldrich | 221473 |
| Epoxy | Devcon | 14250 |
| Biotin-PEG-SVA, MW 5,000 | Laysan Bio Inc. | Biotin-PEG-SVA-5000-100mg |
| mPEG-Succinimidyl Valerate, MW 2,000 | Laysan Bio Inc. | MPEG-SVA-2000-5g |
| TetraSpeck™ Microspheres, 0.1 µm, fluorescent blue/green/orange/dark red | Thermo Fisher Scientific | T7279 |
| Ubp10-mGold2t | Tierra Biosciences | N/A |
| Ubp10(C371A)-mGold2t | Tierra Biosciences | N/A |

